# The Unreasonable Effectiveness of Convolutional Neural Networks in Population Genetic Inference

**DOI:** 10.1101/336073

**Authors:** Lex Flagel, Yaniv Brandvain, Daniel R. Schrider

## Abstract

Population-scale genomic datasets have given researchers incredible amounts of information from which to infer evolutionary histories. Concomitant with this flood of data, theoretical and methodological advances have sought to extract information from genomic sequences to infer demographic events such as population size changes and gene flow among closely related populations/species, construct recombination maps, and uncover loci underlying recent adaptation. To date most methods make use of only one or a few summaries of the input sequences and therefore ignore potentially useful information encoded in the data. The most sophisticated of these approaches involve likelihood calculations, which require theoretical advances for each new problem, and often focus on a single aspect of the data (e.g. only allele frequency information) in the interest of mathematical and computational tractability. Directly interrogating the entirety of the input sequence data in a likelihood-free manner would thus offer a fruitful alternative. Here we accomplish this by representing DNA sequence alignments as images and using a class of deep learning methods called convolutional neural networks (CNNs) to make population genetic inferences from these images. We apply CNNs to a number of evolutionary questions and find that they frequently match or exceed the accuracy of current methods. Importantly, we show that CNNs perform accurate evolutionary model selection and parameter estimation, even on problems that have not received detailed theoretical treatments. Thus, when applied to population genetic alignments, CNN are capable of outperforming expert-derived statistical methods, and offer a new path forward in cases where no likelihood approach exists.

## INTRODUCTION

Using genetic data to make inferences about the natural histories of populations represents a major goal of evolutionary research. As the ever-increasing throughput of DNA sequencing technologies makes the generation of large population genomic data sets more routine, researchers can leverage patterns of genetic variation across the genome to characterize the evolutionary forces at play (Hahn 2018). For example, advances have been made in identifying historical demographic events such as population size changes (Marth *et al.* 2004; Tennessen *et al.* 2012; Gazave *et al.* 2014) and genetic exchange between populations and species (Martin *et al.* 2013; Hellenthal *et al.* 2014; Sankararaman *et al.* 2014; Corbett-Detig and Nielsen 2017; Schrider *et al.* 2018). Population genomic analyses have also revealed the pervasive impact of selection on linked neutral polymorphism (Begun and Aquadro 1992; Begun *et al.* 2007; Langley *et al.* 2012; Elyashiv *et al.* 2016), both through positive selection (Maynard Smith and Haigh 1974; Kaplan *et al.* 1989) and purifying selection (Charlesworth *et al.* 1993). As the volume of population genomic data sets has increased, so too has the demand for powerful computational methods capable of using these data to learn about the fundamental evolutionary processes shaping genomic variation.

To meet this need, myriad statistical and computational tools have been devised to answer evolutionary questions using population genetic data. One particularly common paradigm, which predates the high-throughput sequencing revolution, is that of the population genetic summary statistic: a value (or sometimes a vector of values) designed to capture the information present in a sequence alignment of individuals from one or more populations. When a particular evolutionary phenomenon acts on a population it alters the shapes of genealogies, and this effect is manifest in the observed sequence alignment. For example, a population expansion will result in genealogies with longer branches near the leaves of the tree, which will manifest as an excess of rare alleles. Many summary statistics seek to uncover the signature of these genealogical skews through their effect on the alignment; e.g. Tajima’s *D* will be negative following a recent expansion or recovery from a bottleneck (Tajima 1989; Simonsen *et al.* 1995). Ideally a summary statistic will only detect the signal of the evolutionary process it is being used to investigate, but in practice summary statistics are frequently confounded by other forces that may have similar effects on the shapes and/or sizes of genealogies. For example, Tajima’s *D* is sensitive to positive selection as well as population size changes (Simonsen *et al.* 1995). Moreover, such summary statistics do not capture all of the information present in the alignment. Thus a major challenge of population genetic inference is to create methods that utilize as much information from the input data as possible in order to maximize our ability to distinguish among the numerous evolutionary processes that can give rise to an observed signal.

One approach researchers have adopted to address this challenge is to incorporate a larger number of observations from the data into likelihood-based inference methods. However, calculating likelihoods of population genomic data sets is often mathematically and computationally intractable, and therefore such approaches often use composite likelihoods which ignore the non-independence of observations (e.g. Hudson 2001; Nielsen *et al.* 2005). For example, Nielsen et al.’s SweepFinder (2005), which examines allele frequencies at polymorphisms flanking a focal region to determine whether that region has experienced a recent selective sweep (Maynard Smith and Haigh 1974), treats each allele frequency as an independent observation despite the partially shared evolutionary histories linked alleles experience. Another drawback of most likelihood-based methods is that they generally compute the likelihood of only a few features of the data (often only one), and therefore additional information that could improve accuracy is ignored. For example, SweepFinder examines allele frequencies but ignores linkage disequilibrium (LD), which is elevated in areas flanking the selected site (Kim and Nielsen 2004). Hidden Markov models (Hobolth *et al.* 2007; Boitard *et al.* 2009; Dutheil *et al.* 2009; Kern and Haussler 2010), including those based on the sequential Markov coalescent (Li and Durbin 2011; Schiffels and Durbin 2014), have also proved effective at using population genetic observations along a recombining chromosome to make evolutionary inferences.

More recently, population geneticists have begun to explore an alternative strategy of using a large set of complementary summary statistics for model selection and parameter estimation, an approach that often results in more powerful and robust inference (e.g. Lin *et al.* 2011; Pybus *et al.* 2015; Gao *et al.* 2016; Schrider and Kern 2016; Sheehan and Song 2016). Each summary statistic seeks to measure a particular attribute of the genealogy, and one can thus design a customized set of summary statistics to more fully represent the genealogical information present in the sequence alignment. This view deploys summary statistics less for their individual links to underlying theory, and more for their collective ability to perform pattern recognition. The challenge then becomes extracting information about the underlying evolutionary processes from the set of summary statistics. Two exciting approaches for dealing with this challenge that have garnered increasing attention in recent years are approximate Bayesian computation (ABC; reviewed in Beaumont 2010) and supervised machine learning (reviewed in Schrider and Kern 2018). Both of these approaches make use of suites of user-defined summary statistics and training data generated under known parameters to identify reasonable evolutionary models and parameterizations that could have generated the observed data. Here we focus on the supervised machine learning approach, as it sets the scene for the convolutional neural networks described below.

In the terminology of supervised machine learning, each summary statistic is called a feature, and the full set of statistics used is called a feature vector. To use supervised machine learning, a researcher must first obtain training data (often referred to as “labeled” data)—a set of data points each summarized by a feature vector (the explanatory variables) accompanied by a known outcome (the response variable). Next, a supervised machine learning algorithm is trained to predict the outcome given the feature vector using the labeled training data. Thus, the supervised machine learning technique automates the process of extracting information and constructing rules from a set of summary statistics. Across many areas of research, supervised machine learning techniques are fast replacing rules developed by human experts because they are often more accurate (LeCun *et al.* 2015).

Supervised machine learning methods are increasingly being applied to numerous problems in population genetics (Schrider and Kern 2018). In this context, labeled training data are usually generated via population genetic simulation, an endeavor that has grown considerably more feasible given recent improvements in simulation flexibility and efficiency (e.g. Thornton 2014; Kelleher *et al.* 2016; Haller and Messer 2017; Kelleher *et al.* 2018). To date, population genetic applications of machine learning include demographic inference (Pudlo *et al.* 2016; Sheehan and Song 2016), local ancestry inference (Schrider *et al.* 2018), inferring recombination rates (Lin *et al.* 2013; Gao *et al.* 2016), and detecting genomic regions experiencing recent selective sweeps (Pavlidis *et al.* 2010; Lin *et al.* 2011; Ronen *et al.* 2013; Pybus *et al.* 2015; Schrider and Kern 2016). While such methods have great promise, they still rely on a user-defined set of summary statistics (ranging in number from dozens to hundreds). Moreover, it is not known whether it is possible to construct a set of statistics that sufficiently captures all relevant information in the input data.

Unlike other machine learning approaches, convolutional neural networks (CNN; LeCun *et al.* 1998) are pattern recognition algorithms that do not require a predefined feature vector. When fed labeled training data (e.g. a set of haplotypes simulated under a known biological scenario), a CNN discovers meaningful features, in essence making a feature vector, and then extracts information from these features in order to make inferences. CNNs have proved effective in a number of fields (reviewed in LeCun *et al.* 2015), and particularly in the field of image recognition, where they have achieved dramatic improvements over previous efforts (e.g. Lawrence *et al.* 1997; Krizhevsky *et al.* 2012; Simonyan and Zisserman 2014). The application of CNNs to population genomic inference is just beginning, and shows great promise (Chan *et al.* 2018). Population genetic questions may be particularly well suited for CNN-based learning because they take matrices as inputs, and alignments of sequenced chromosomes are quite naturally represented in this manner.

The goal of this paper is to assess the effectiveness of CNNs as a general strategy for population genomic inference. We demonstrate that CNNs can be successfully applied to a number of population genomic problems, in some cases achieving surprising accuracy. In particular, we use simulation to show that CNNs can leverage images of aligned sequences to accurately uncover regions experiencing gene flow between related populations/species, estimate recombination rates, detect selective sweeps, and make demographic inferences. Indeed, in most cases we observe performance that matches or exceeds that of current methods. We also use a CNN to accurately infer recombination rates from read coverage data in a simulated autotetraploid, demonstrating this approach’s flexibility in handling noisy data while solving a complex problem for which no theoretical solution exists. In light of these encouraging findings, we argue that population genetics researchers should consider CNNs as a potential solution to a variety of problems involving evolutionary inferences from sequence data. Because some readers may have little background with this tool, we also provide an overview of the inner workings of CNNs and explore several technical considerations that may impact performance.

## RESULTS

Our goal is to use a CNN to make population genetic inferences from an alignment image, which can be thought of as matrices where each entry represents the allele present in a given chromosome at a given site. In particular, we focus on four distinct problems: identifying local introgression, estimating the recombination rate, detecting selective sweeps, and inferring population size changes. We chose these four tasks because each represents a different challenge in population genetic inference, each with its own attendant branch of theory. To show the ability of CNNs to solve problems for which no statistical approaches have been proposed, we extended our recombination inference to infer recombination rates in autotetraploids with tetrasomic inheritance.

Below, we address each of these problems in turn, providing a brief overview of the phenomenon in question and existing methodology before describing our results using CNNs. But prior to tackling these problems, we first give an overview of CNNs and discuss strategies for reorganizing our input data that we found helpful in making CNNs work more efficiently with population genetic alignments.

### Overview of convolutional neural networks

Internally, a CNN is a type of artificial neural network – a collection of connected layers of combinatorially linked mathematical functions (termed *artificial neurons*) that take an input and transform it into an output value (Mitchell 1997). In a typical fully connected artificial neural network, the input values are fed through a series of layers of artificial neurons (fig. 1A), termed hidden layers, before reaching the output layer which transforms its inputs into a final prediction. The output for the *j*^th^ neuron within one of the hidden layers is given by the following:

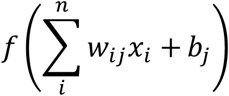

In the expression above, *x*_*i*_ is the neuron’s *i*^th^ input value (either an input value from the data or from a neuron in the previous layer’s output), and *w*_*ij*_ is the *weight* attached to the connection between that node (*i*) and the current node (*j*) and *b*_*j*_ is the current node’s *bias* term. That is, to obtain the value of neuron *j*, we compute the linear combination of the vector containing all values from the previous layer and the *j*^th^ neuron’s vector of weights; the results of this summation are in turn added to neuron *j*’s bias term and then fed as input to some function *f*, termed the *activation function* and which may be nonlinear. Thus, an artificial neural network is a mathematical function.

**Fig. 1:**
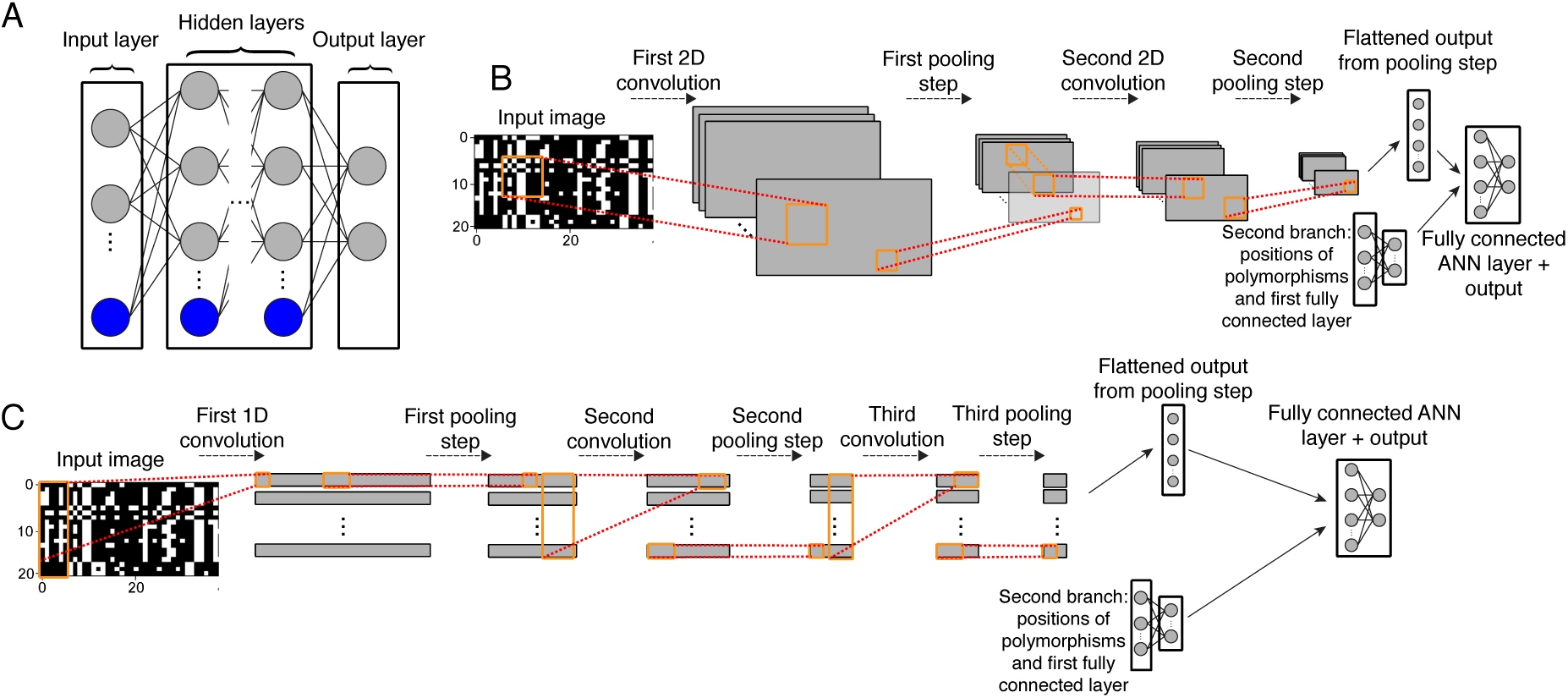
Schematics of a standard feedforward neural network and two convolutional neural network designs used in this study. A) Diagram of a fully connected feedforward neural network. Gray circles represent input (left side), output (right side), or hidden (center) neurons. Blue circles represent collections of bias terms. With the exception of the input layer, the value of any given neuron is a linear combination of values from the previous layer plus a bias term; this sum is then passed to an activation function (not shown). Each edge represents a distinct weighted input or bias term. Outputs may represent class membership posterior probabilities or estimates of continuous variables. B) A diagram of a 2D CNN similar to that used in this study to infer demographic parameters. The input is an alignment represented as an image which is passed through a first convolutional layer in order to create a set of feature maps. These feature maps are then downsized via a pooling step which replaces the values of a 1 or 2D matrix within a feature map with a single value summarizing it (e.g. the mean or maximum value of that matrix). For example, a 2D pooling operation of size 2 will reduce the size of a feature map by a factor of 4, as each adjacent 2×2 matrix within the input feature map is replaced by a single value (e.g. the maximum of those four values). These downsized feature maps are then passed through a second convolutional filter and pooling step, and the resulting output is flattened into a one dimensional vector and passed as input into a fully connected feedfoward layer (bias terms not shown). Also passed into this layer is output from a second branch of this network: the vector of positions of segregating sites in the alignment which have been passed through their own fully connected layer. Finally, the last fully connected neural network layer yields the predicted output values. C) Similar to panel B, but showing a 1D CNN with three convolutional layers (each followed by a pooling step), as used for our recombination rate estimator.

Importantly, by changing the values of the weights and biases, an artificial neural network can be tuned to detect informative patterns in the input data in order to produce the desired output. In the case of image recognition, an image is first represented numerically, typically as a matrix of pixel intensities, and then transformed by the artificial neural network to produce an output, for example a prediction of the type of object in the image. CNNs (Fig. 1B–C) differ from standard artificial neural networks in that they begin with one or more convolutional layers, in which a series of smaller weight matrices referred to as “filters” slide across the input image—mimicking the manner in which animal cortical neurons each focus on input only from a small receptive field—and perform a matrix convolution at each step until a series of filtered image matrices are produced (LeCun *et al.* 1998). These filters are constructed during training (see below). Each convolutional layer is often followed by a pooling layer (see Fig 1B and caption) which reduces the size of these filtered image matrices while maintaining potentially important discriminatory information obtained by the convolutional filters. Finally, these matrices are flattened into one-dimensional vectors and then fed into a fully connected (or “dense”) artificial neural network (for an accessible overview see LeCun *et al.* 2015). Thus, salient features derived from the image matrix by the convolutional and pooling layers are passed into one or more layers of a fully connected neural network whose output layer then yields our predicted response value.

CNNs allow for two types of convolutional layers: 1-dimensional and 2-dimensional, which differ only with respect to the possible shapes that the convolutional filter can take (Fig. 1B–C). 1-dimensional (1D) convolutions are often used in the application to time-series data (e.g. Dieleman and Schrauwen 2014; Kim 2014), and are thus applicable to sequence alignment matrices. Despite its name, a 1D filter is not a vector but rather a rectangular matrix that spans a user-defined number of entries (called the “kernel size”) in one dimension in the input data (in our case this dimension is that of the polymorphic sites in the alignment), and stretches entirely across the other dimension (in our case across all chromosomes in the sample). A 2-dimensional (2D) convolutional filter, which is more often used with image data, allows the user to specify both dimensions of the filter matrix (often using a square matrix). Whether 1- or 2-dimensional, the benefit of incorporating convolutions is that it allows the CNN to take advantage of structural information in the input data. For example, from an image of a face, a CNN can learn to detect the repeated pattern of the eye shape and the location of both eyes relative to one another and to other features. When there is meaningful structural information such as this, CNNs tend to outperform non-convolutional neural networks.

Here our input data is an alignment of linked segregating sites with partially shared evolutionary histories. Our hope is that a CNN can discover structural information in these data in order to make evolutionary inferences—for example, locating the valley in diversity at the center of a sweep (Maynard Smith and Haigh 1974), the “shoulders” on the flanks of a sweep where linkage disequilibrium and allele frequencies are both elevated (Schrider *et al.* 2015), or even the spatial relationship between these patterns. We also note that neural networks such as CNNs can have multiple “branches” each with separate architectures and input types—in some of the cases discussed in this paper we incorporate an additional network branch whose input is the vector of the positions of the segregating sites (Fig. 1B–C).

Like all supervised machine learning methods, a CNN must be trained on labeled training data before it can make predictions on unlabeled data (i.e. data whose response variables are unknown). Training is accomplished by tuning the weights and biases that control the behavior of its artificial neurons so that together they maximize the accuracy of the outputs on the training data. Note that the weights determined during the training process include the values of the convolutional filter matrices, and thus different filters will be algorithmically created for each task we address in this paper. This tuning occurs over a number of iterations using the backpropagation algorithm (Rumelhart *et al.* 1986), which in modern implementations feeds a small number of training examples (a “mini-batch”) through the network and then estimates the error gradient on the output vectors produced for these examples. The error gradient is then propagated in reverse through the network—a given hidden neuron’s contribution to the error is proportional to the linear combination of its weight vector and the errors associated with each neuron in the next layer. The weights are then updated using one of the many flavors of stochastic gradient descent (e.g. Kingma and Ba 2014). This process repeats until each training example has been fed through the network, marking the completion of a single training iteration. Training continues for a number of these iterations (often called epochs) until a specified stopping criterion is reached (e.g. a predefined number of iterations has been performed, accuracy on the validation set has not improved relative to the previous iteration, etc.).

In the context of population genetics, the CNN’s input could be a matrix of allelic states at each polymorphic site (Fig. 2). For example, an alignment of haploid individuals *M*, where *M*_*ij*_=0 if the *i*^th^ individual has the ancestral allele at the *j*^th^ segregating site in the alignment, and 1 if this individual has the derived allele (an input format that can easily be altered to allow for multiallelic polymorphisms); we adopt this approach and variants of it below. The output can be a categorical indicator (e.g. whether or not the genomic window experienced a recent selective sweep) in which the problem is referred to as a classification task in machine learning terminology, a quantitative value (e.g. the population recombination rate) in which case the task is referred to as regression, or a vector containing both categorical and quantitative values. Once the CNN has been trained to produce the desired output, it can be applied to unlabeled data (e.g. sequence from natural populations).

**Fig. 2:**
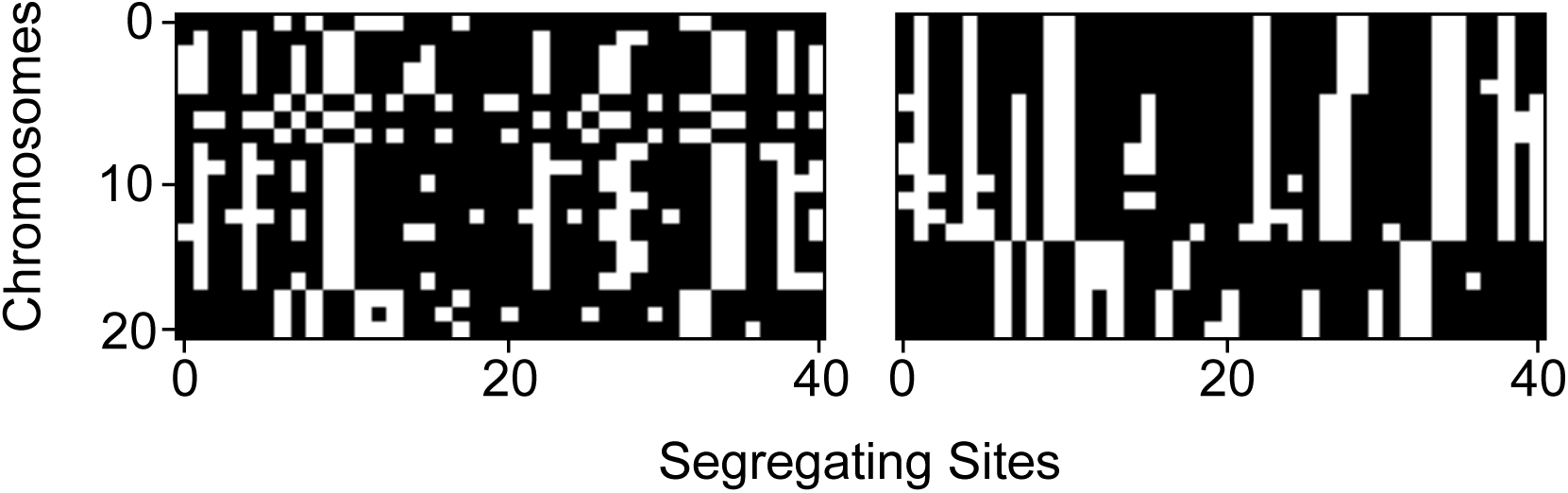
Example population genetic alignments visualized as black-and-white images. An unsorted alignment matrix (left) and this same matrix sorted by genetic similarity among chromosomes (right) are shown. Each row represents one of twenty chromosomes in the sample and each column represents one of forty segregating sites. Derived and ancestral states are encoded as black and white, respectively.

Because supervised machine learning relies on predictive functions tuned algorithmically from training data, CNNs can be applied to any problem for which a training set can be obtained, and therefore our inference is not limited to problems for which appropriate likelihood models or statistics have been derived and implemented. In a population genetics context, coalescent simulations provide a versatile and computationally efficient (Hudson 2002; Teshima and Innan 2009; Ewing and Hermisson 2010; Kelleher *et al.* 2016; Kern and Schrider 2016) means to generate training data. In this paper we relied exclusively on coalescent simulations to produce training data for the CNN. However, compute-intensive forward population simulations may offer greater flexibility than coalescent simulations in some situations, and recent advances are making them more computationally feasible (Kelleher *et al.* 2018).

### Using a CNN to make inferences from an alignment: a simple test case

We evaluated the performance impact of transposing the alignment matrix (so that columns rather than rows correspond to chromosomes) and sorting the chromosomes in the alignment matrix by genetic similarity. We did this using a 1D CNN trained to estimate the population-scaled mutation rate, *θ*, in an equilibrium population. We found that both of these techniques accelerate the decline in root-mean-square error (RMSE; Fig. 3), showing that they help the network achieve better performance. Transposing the alignment matrix so that chromosomes are represented by rows and polymorphisms by columns has a particularly notable effect (compare blue and black lines in Fig. 3). Additionally, sorting the chromosomes by genetic similarity further increases the accuracy of the CNN when combined with the matrix transposition above (magenta line); alternatively, using a permutation-invariant network architecture would obviate any need for this step (Chan *et al.* 2018). The effect of transposition should disappear when using 2D convolutions because in those cases we always used a square convolutional filter matrix (Methods), but we found that 1D CNNs often performed as well as 2D CNNs (data not shown). Thus, unless otherwise specified we use 1D convolutions for the tasks discussed below.

**Fig. 3:**
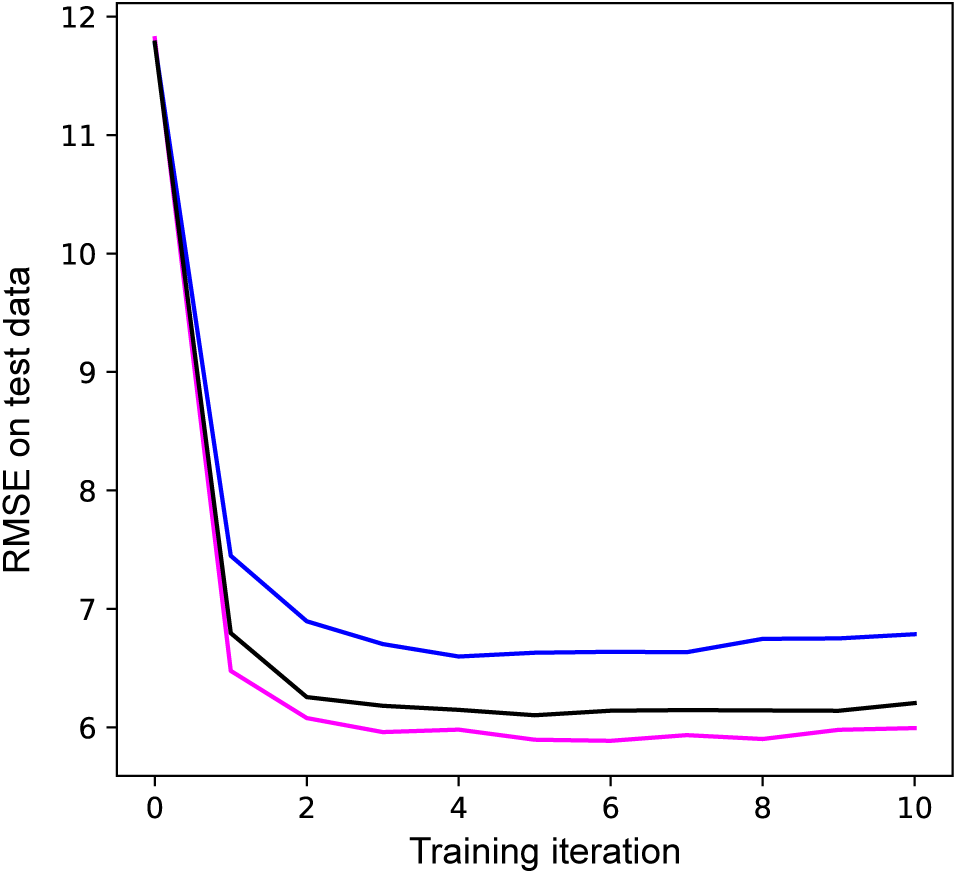
The impact of input data reorganization on accuracy. We show the root mean squared error (RMSE) of a 1D CNN’s predictions of *θ* as assessed on 1,000 test alignments after a given number of training iterations. Each line is the average of 10 runs. The blue line shows accuracy after training using alignment matrices with each row representing one chromosome. The black line shows accuracy after transposing all matrices so that chromosomes correspond to columns; this makes 1D convolutional filters examine each individual at a group of adjacent segregating sites. The magenta line shows the impact of transposing matrices, and sorting the chromosomes in the alignment matrix by genetic similarity.

### CNN’s can accurately detect introgressed loci

Recent studies indicate that closely related species often exchange genes (Kulathinal *et al.* 2009; Martin *et al.* 2013; Brandvain *et al.* 2014; Fontaine *et al.* 2015). There are several motivations for locating genomic segments introgressed from one species into another. For one, the occurrence of cross-species gene flow raises the possibility of adaptive introgression, wherein a beneficial allele enters a population via migration from a related species (reviewed in Hedrick 2013). Discovering introgressed loci can therefore identify alleles underlying rapid ecological adaptation as well as the source of these alleles. In addition, uncovering genomic regions that are and are not porous to cross species gene flow may help to illuminate the genomic basis of reproductive isolation (Turner *et al.* 2005).

Researchers have thus sought to devise methods capable of detecting introgressed regions from multispecies population genomic data sets. These include methods that attempt to infer the ancestry for each individual at each site (e.g. Price *et al.* 2009; Lawson *et al.* 2012; Sohn *et al.* 2012) and those that explicitly seek to discriminate between introgressed and non-introgressed loci (Sankararaman *et al.* 2014; Geneva *et al.* 2015; Rosenzweig *et al.* 2016; Schrider *et al.* 2018). We trained a CNN to identify introgression in a scenario modeled after the demographic history of the *Drosophila simulans*-*D. sechellia* species pair (Methods), for which there is evidence for recent gene flow (Garrigan *et al.* 2012).

Fig. 4A displays the results of these tests in the form of confusion matrices, which show the fraction of test examples correctly predicted for each class (diagonal values) as well as the fractions incorrectly assigned (off-diagonal values). To compare the performance of our CNN to competing approaches, Fig. 4B displays the confusion matrix for FILET, a method previously shown to outperform several methods, including two statistics for detecting introgression (Joly *et al.* 2009; Geneva *et al.* 2015), and a tool that infers local ancestry tracks for each individual (Lawson *et al.* 2012). Overall, this CNN classified 88.5% of test simulations correctly (95% confidence interval: 87.7–89.2%). The most difficult scenario for the CNN was introgression from *D. simulans* into *D. sechellia*, which it misclassified as “no introgression” 23% of the time. For the other two classes the CNN accuracy was >95%. Importantly, for every class this CNN achieved greater accuracy than FILET (overall accuracy of 82.5%; 95% confidence interval: 81.7%–83.4%), a machine learning approach that leverages a vector of 31 summary statistics (Schrider *et al.* 2018). Thus, it is a useful measuring stick for assessing the CNN’s accuracy, and the CNN’s success in this comparison is encouraging.

**Fig. 4:**
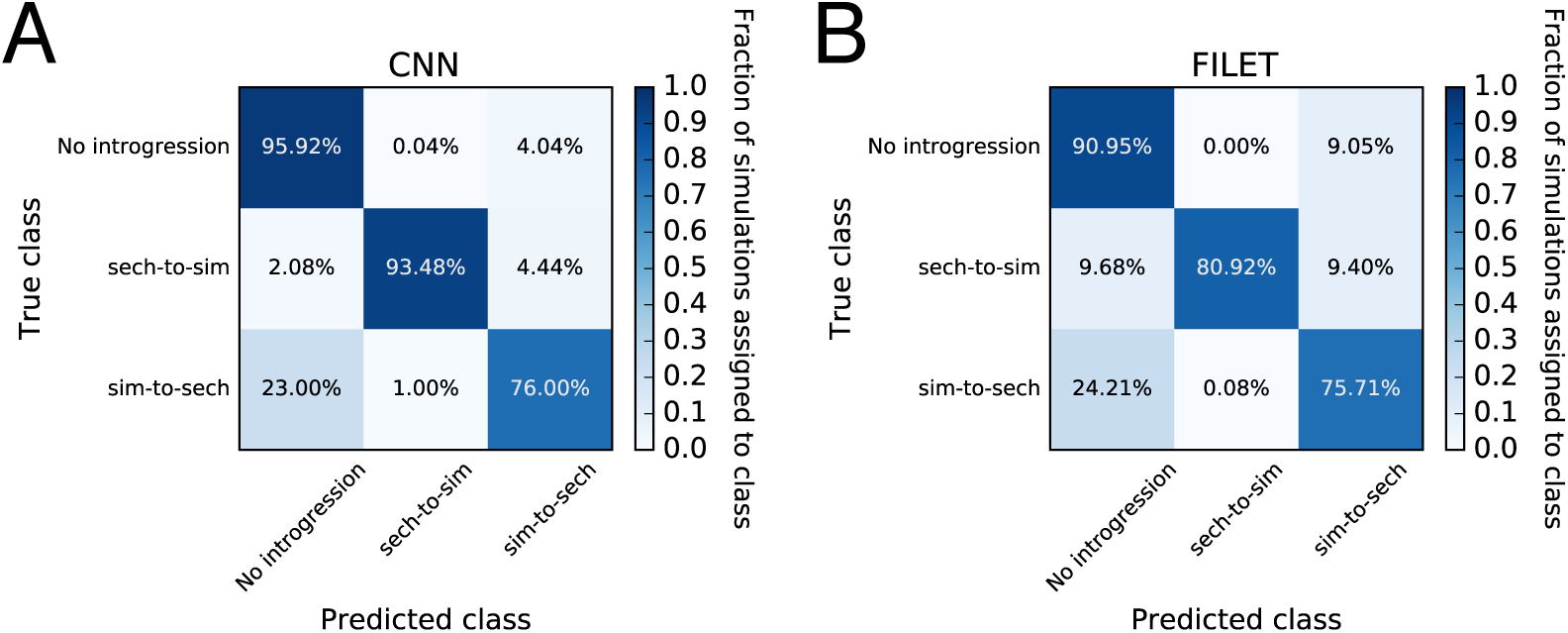
Performance of classifiers for detecting introgression. We use confusion matrices to show the performance of a CNN trained to detect genomic regions of introgression between two closely related species (panel A), and a competing method that uses a vector of summary statistics to the same end (FILET; panel B). These classifiers were both trained and tested on the same data sets which were simulated under a joint demographic history inferred from a sample of *Drosophila simulans* and *D. sechellia* individuals (as described in the Methods) with and without introgression. The classifiers seek to discriminate among three classes: no introgression in the genomic window being examined, introgression from *D. sechellia* to *D. simulans*, and introgression from *D. simulans* into *D. sechellia*. Each entry in the matrix shows the fraction of test examples belonging to the class specified on the *y*-axis that were inferred by the method to belong to the class specified on the *x*-axis. Correct classifications are those found along the diagonals, while all off-diagonal entries represent incorrect classifications.

### Estimating historical recombination rates

Recombination creates new combinations of alleles, and the degree of linkage between selected sites affects the efficiency with which natural selection can act on each individual site (Hill and Robertson 1966). The interplay of selection and recombination also influences the landscape of diversity across the genome (Begun and Aquadro 1992). Knowledge of recombination rates is thus key to population genetics research. As a more practical alternative to estimating rates directly (e.g. from pedigrees; Kong *et al.* 2010), one can infer recombination rates from population genetic data by examining associations among alleles at different sites. A number of methods have been proposed to solve this problem, including summary statistic estimation approaches (e.g. Hudson and Kaplan 1985; Hudson 1987; Hey and Wakeley 1997), composite likelihood-based methods (e.g. Hudson 2001; McVean *et al.* 2004; Chan *et al.* 2012), and machine learning tools using a vector of statistics (Lin *et al.* 2013; Gao *et al.* 2016). We sought to determine whether a CNN taking an alignment image as input could be trained to tackle this task. To address this problem, we first trained a CNN to estimate the historical population recombination rate 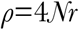 (where *r* is the crossover rate per base pair per meiosis) from phased chromosomes. This is the simplest scenario, as the arrangement of alleles on chromosomes is completely resolved. Following training, we compared the CNN’s performance to that of LDhat (McVean *et al.* 2004), a widely used composite likelihood method, on the same testing data (Fig. 5). We generated a test set of alignments whose values of *ρ* spanned three orders of magnitude, from 0.0002 to 0.2 (expressed per bp). Overall, both approaches performed well at predicting the true value of *ρ*. LDhat had an *R*^2^ = 0.77 and an RMSE = 0.016, whereas the CNN had a *R*^2^ = 0.86 and an RMSE = 0.011 (Fig. 5A,B). LDhat appears to estimate *ρ* slightly better than the CNN for lower recombination rates, whereas the CNN performs better at the higher values of *ρ* (Fig. 5C). Additionally, the CNN appears to provide a roughly unbiased estimator of *ρ*, while LDhat’s estimates appear downwardly biased.

**Fig. 5:**
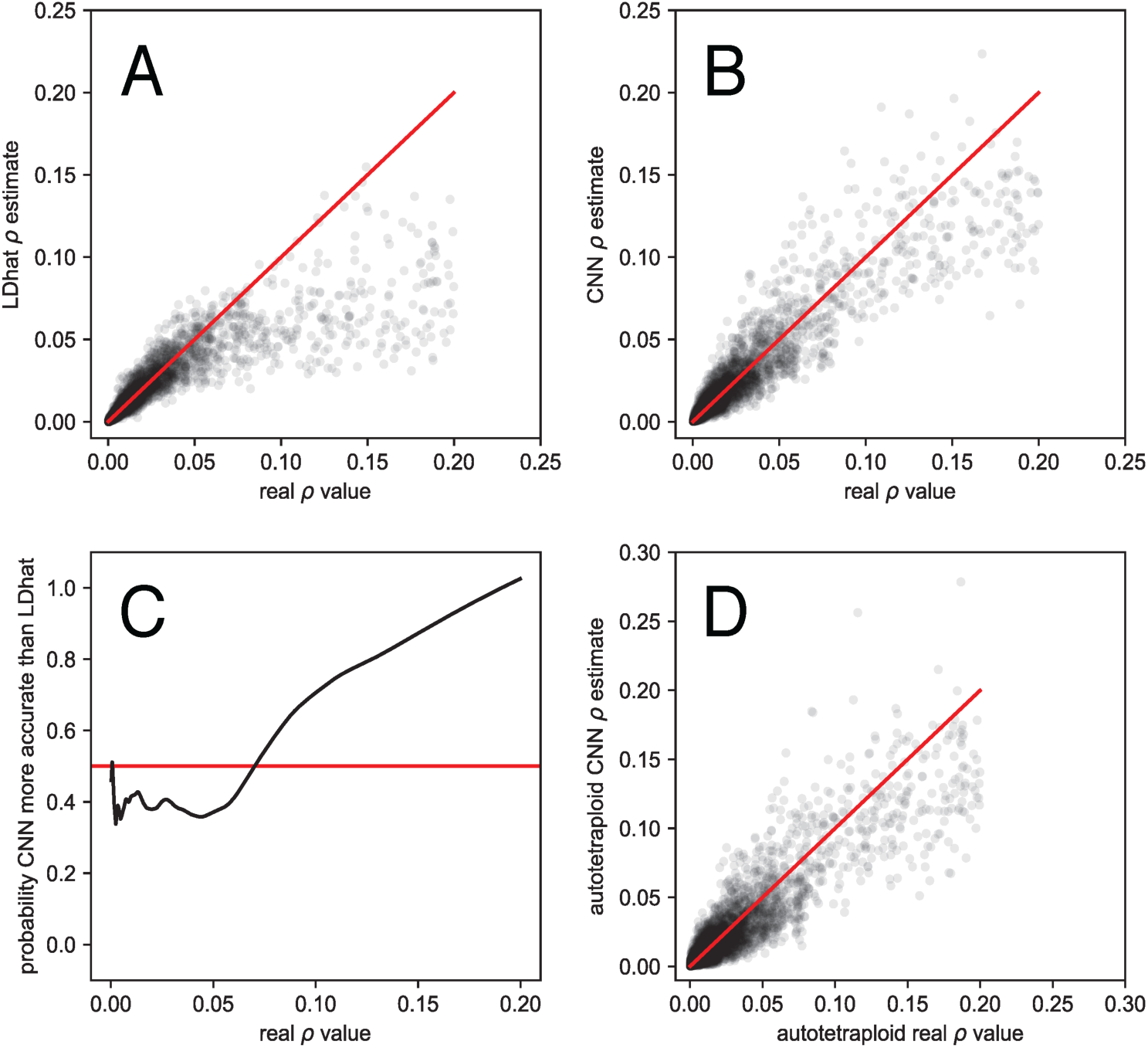
Accuracy of recombination rate estimates from LDhat and our CNN. Panels A and B show the real *ρ* values per base pair on the *x*-axes and LDhat’s (A) and the CNN’s (B) predictions on the *y*-axes. Panel C again shows the real *ρ* values on the *x*-axis, and the probability that the CNN was more accurate than LDhat (black line) on the *y*-axis. This probability was calculated by scoring estimates where the CNN outperformed LDhat as one and the reciprocal as zero, and then smoothing these values with a lowess curve with a span of 15%. The red line represents the expectation if both methods had identical accuracy. Panel D shows the results from the simulated autotraploid model, with the real *ρ* values on the *x*-axes and the CNN prediction on the *y*-axes.

Because the CNN was capable of estimating *ρ* independent of *θ*, we were interested to see how well it could interpolate between the *θ* values it was trained with. The CNN was trained with a large gap between *N* = 20,000 and *N* = 50,000 (and thus a large gap in *θ*; see Methods), so we used coalescent simulations to generate an additional test set with *N* values drawn uniformly among 30,000, 35,000, 40,000, and 45,000. When tested on these data the CNN’s predictions had an *R*^2^ = 0.82 and an RMSE = 0.017. This represents a slight decrease in accuracy from the values obtained when tested on the same *N* values used in training, but nonetheless shows that the CNN can interpolate between training parameters without a dramatic loss in accuracy. This could be a useful property, for example in cases where *N* (or *θ*) is unknown, but where one can generate coalescent simulations across a range of plausible values.

Further complications arise when estimating *ρ* from unphased data. Under this scenario the arrangement of alleles on chromosomes is not known. One work-around is to first phase the alleles and then infer *ρ* as above, but not all data sources are easily phased, and phasing errors will, of course, reduce accuracy. Another approach is to analyze the unphased data directly. The relevant theory required to tackle this problem in a probabilistic manner has been worked out for unphased diploids (Auton and McVean 2007), but expanding this theory to higher ploidies would require a substantial effort. Take for example an autotetraploid with tetrasomic inheritance, where there are five possible genotypes (*AAAA*, *AAAa*, *AAaa*, *Aaaa*, and *aaaa*). To further complicate things, after sequencing an autotetraploid genome to a moderate depth of coverage and identifying polymorphisms, the true underlying genotype may be uncertain. For example, given a site with 10 reads supporting *A* and 10 supporting *a*, the true genotype could be *AAAa*, *AAaa*, or *Aaaa*. To show the utility of CNNs in addressing novel population genomic inference problems, we designed a CNN capable of inferring *ρ* from a simulated set of sequence reads from an unphased autotetraploid population sample.

We used a simple simulation scheme to produce read counts for each allele at each site for each individual in a sample of 12 autotetraploids, each with approximately 25X expected genome-wide coverage (see Methods). Rather than allelic assignments, the input matrix for this CNN contains for every site in each individual the fraction of reads bearing the *a* allele. Deriving a likelihood function for *ρ* under this formulation may be challenging, and such a solution has not yet been attempted. However, appropriately designed artificial neural networks are universal approximators, meaning that they have the potential to approximate any continuous function over a compact input space (Hornik 1991). Thus it is possible for a CNN to approximate the desired likelihood function, even in its absence. To this end we trained a CNN with a similar architecture to the one used above on phased haploid chromosomes (see Methods). We evaluated the performance of this CNN on a set of simulations where *ρ* again ranged from 0.0002 to 0.2 (still scaling by 4*N*, rather than 8*N* which would be appropriate for tetraploids, so the result can be compared to those above). The CNN’s predictions had an *R*^2^ = 0.83 and an RMSE = 0.012 (Fig. 5D). As before, the estimate of *ρ* was made independent of *θ*, which varied over an order of magnitude. The fact that this autotetraploid network performed only slightly worse than the haploid version demonstrates that a CNN can solve problems for which no model-based likelihood (or even composite likelihood) approach has been obtained, empowering empiricists untrained in methods development to address questions specific to their biological system.

### CNNs can accurately detect and categorize signatures of recent positive selection

When a new mutation is immediately favored by positive selection, it rapidly increases in frequency until it fixes (i.e. completely replaces all other alleles at that site). This phenomenon, referred to as a hard selective sweep, drastically reduces the amount of linked neutral variation (Maynard Smith and Haigh 1974), and produces characteristic skews in the allele frequency spectrum (Fay and Wu 2000) and linkage disequilibrium at linked sites (Kim and Nielsen 2004). Alternatively, in a process known as a “soft sweep” populations may adapt via selection on a polymorphism that has been segregating for some time, such that the adaptive allele exists on numerous haplotypes (Hermisson and Pennings 2005). To uncover the mode of recent adaptation and the genomic regions underlying recent adaptation, a large number of methods have been devised to detect and characterize selective sweeps. These include summary statistics (Kelly 1997; Fay and Wu 2000; Kim and Nielsen 2004; Voight *et al.* 2006; Garud *et al.* 2015), composite likelihood-based approaches (Kim and Stephan 2002; Kim and Nielsen 2004; Nielsen *et al.* 2005; Vy and Kim 2015), and supervised machine learning approaches using a vector of statistics to obtain greater power than individual tests/statistics (Lin *et al.* 2011; Pybus *et al.* 2015; Schrider and Kern 2016; Sheehan and Song 2016; Sugden *et al.* 2018). Although these efforts have led to considerable progress, detecting and distinguishing between hard and soft sweeps remains a major challenge.

We built a CNN to detect selective sweeps and to discriminate between hard sweeps and soft sweeps. This CNN follows the S/HIC method of Schrider and Kern (2016) by casting the problem as a classification task where the genomic region being examined is assigned to one of five disjoint classes: a recent classic “hard” sweep, a recent “soft” sweep, a region linked to a nearby hard sweep, a region linked to a nearby soft sweep, or a neutrally evolving region.

Like FILET for the problem of detecting introgression, comparing the CNN’s accuracy to that of S/HIC is informative because S/HIC was previously shown under a variety of simulated scenarios to have greater power than a number of competing methods (Schrider and Kern 2016). Rather than adopting S/HIC’s approach of using a large vector of statistics, the CNN takes an alignment image as input. We tested both methods against data simulated under a challenging demographic history estimated from human population data (Methods). As evidenced by the confusion matrices in Fig. 6, the CNN has slightly higher overall accuracy than S/HIC (60.6% with 95% confidence interval: 58.8–62.3% for the CNN; versus 58.5% with 95% confidence interval: 56.7%–60.2% for S/HIC). While S/HIC appears to be somewhat more sensitive to sweeps, the CNN is achieves a more than 3-fold reduction in false positive rate: 2% of neutral simulations are classified as sweeps by the CNN, versus 6.35% for S/HIC; all of these false positives are classified as soft sweeps. This quality may be particularly desirable when scanning genomes where sweeps are relatively rare and thus a high degree of specificity is required to maintain a low false discovery rate, although the proclivity of either classifier to produce false positives versus false negatives can be adjusted by imposing a posterior probability cutoff. Note that these classifiers were both trained under the same demographic history from which the test data were generated. We would not expect this CNN to match S/HIC’s robustness to demographic misspecification given that S/HIC’s feature vector was designed with this in mind, though we did not test this. Nonetheless, the fact that the CNN has similar accuracy to S/HIC under this difficult test scenario is highly encouraging.

**Fig. 6:**
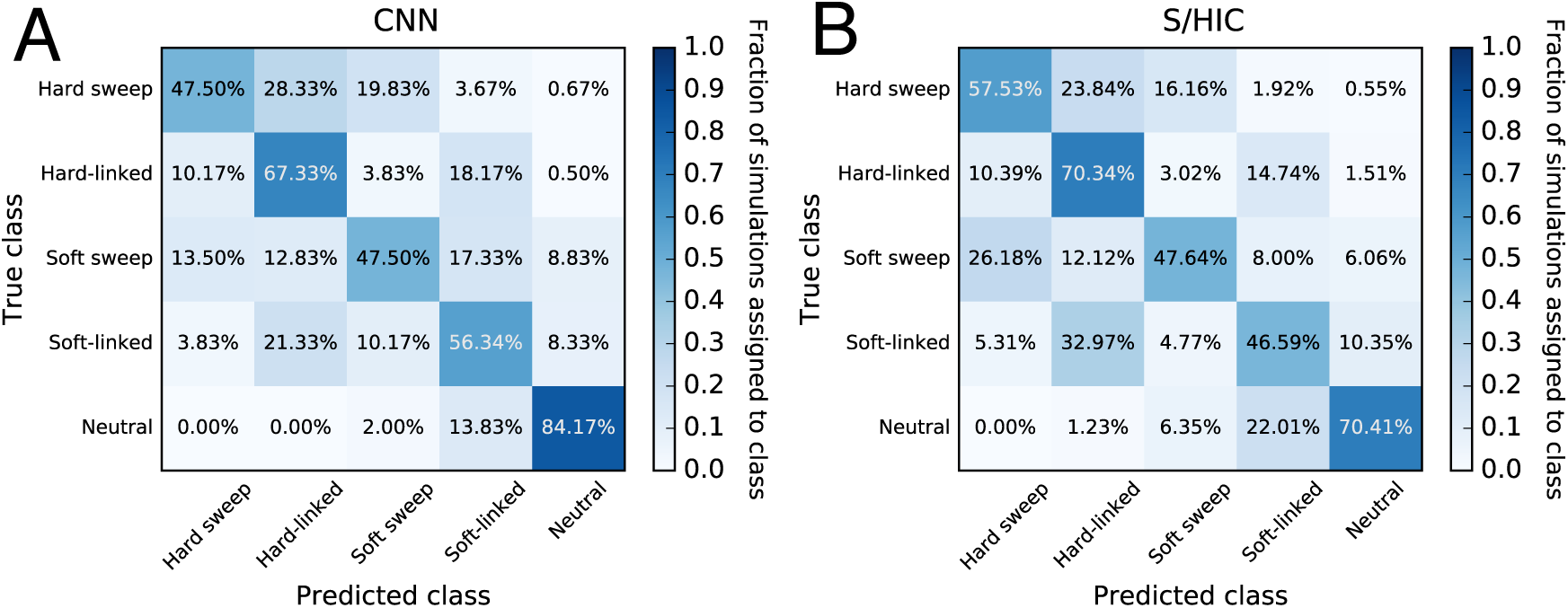
Confusion matrices showing accuracies of two methods that seek to detect recent positive selection by discriminating among hard sweeps, soft sweeps, unselected regions closely linked to hard and soft sweeps, and neutrally evolving regions. (A) Confusion matrix summarizing the performance of our CNN, which uses an alignment image as input. (B) Performance of S/HIC, which uses a vector of summary statistics each measured in windows surrounding the region to be classified. These two classifiers were both trained and tested on the same data sets described in the Methods.

### CNNs can extract demographic information from alignments

A major focus of population genetics research is to use genomic data to elucidate species’ demographic histories—the extent and timing of population size changes, and the history of population splits and migration events. For example, a host of population genetic approaches have been devised to infer the times and intensities of population contractions and expansions over the course of a species’ recent history (e.g. Marth *et al.* 2004; Schiffels and Durbin 2014; Liu and Fu 2015), and to elucidate the history of population splits and subsequent gene flow (Nielsen and Wakeley 2001; Hey 2009), and population merging events (e.g. Lipson *et al.* 2013; Loh *et al.* 2013). We asked whether CNNs can effectively extract demographic information from alignment images, focusing on the task of inferring population size histories. In particular, we attempted to train a CNN to estimate the parameters of a three-epoch model of instantaneous effective population size changes. There are five such parameters: the ancestral population size (*N*_*2*_), the time of the more ancient population size change (*T*_*2*_), the population size after this change (*N*_*1*_), the time of the more recent change (*T*_*1*_), and the present-day population size (*N*_*0*_); our response variable is the vector of these 5 real-valued parameters. Thus this analysis also allows us to assess the ability of CNNs to predict multiple population parameters simultaneously.

We simulated 50 haploid chromosomes under a variety of randomly selected population size histories, and trained a CNN to estimate the demographic model parameters. The simulated region was roughly equivalent in length to 1.5 Mbp of the human genome (Methods). Because we found this problem to be comparatively difficult, we experimented with a variety of hyperparameters governing the neural network structure and input/output format. In supplementary table S1 we show the optimal RMSE (i.e. the minimum RMSE across training iterations) for each hyperparameter combination examined. This experiment revealed several general trends. First, 1D convolutional networks tended to fare slightly better than their 2D counterparts (median RMSE of 0.52 across all hyperparameter combinations with 1D convolutional filters, and median RMSE 0.54 for 2D convolutions; *p*=1.1×10^-4^; Mann-Whitney *U* test); however several 2D networks performed nearly as well as the best 1D network, achieving an RMSE of <0.5 while the best score obtained overall was 0.43. Second, smaller convolutional filters tended to perform slightly better than larger ones—we observed a positive correlation of kernel size with RMSE across hyperparameter combinations (*ρ*=0.26; *p*=6.9×10^-4^; Mann-Whitney *U* test). For example, the median validation RMSE was 0.51 for a kernel size of 2 versus 0.56 for a kernel size of 10. Third, log-scaling the demographic parameters to be estimated increased accuracy (RMSE decreased from 0.55 to 0.52; *p*=0.020; Mann-Whitney *U* test). For this problem sorting chromosomes by relatedness resulted in a small improvement (RMSE decreased from 0.54 to 0.53; *p*=0.034). Encoding ancestral and derived alleles as ‘0’ and ‘255’ (i.e. black and white in a grayscale image), respectively, versus ‘-1’ and ‘1’ had a significant influence on accuracy, with the former yielding better performance than the latter (RMSE of 0.51 vs. 0.60; *p*=1.5×10^-15^). Finally, using dropout resulted in a slight decrease in accuracy (median RMSE increased from 0.52 to 0.55) though this was not statistically significant (*p*=0.092). We note that these trends may change if the amount of training data is increased or decreased, and may not necessarily hold for other tasks.

In Fig. 7, we show the correlation between the true and inferred values for each of these 5 parameters for the best performing network. For *N*_*0*_ and *T*_*0*_, these correlations are quite high, implying that our CNN can recover the true values reasonably well. However, for the remaining parameters, the correlation is lower (though still highly significant), and our CNN produces downwardly biased estimates when the values of these parameters are larger. Although our accuracy is far from perfect, we consider these results fairly encouraging because we are only examining a single moderately sized genomic region, while other modern demographic inference methods use data from across the genome. For example, ∂a∂i (Gutenkunst *et al.* 2009) uses allele frequencies measured at a large number of polymorphisms (e.g. those found in all distal intergenic regions across the genome; Gazave *et al.* 2014). PSMC and MSMC (Li and Durbin 2011; Schiffels and Durbin 2014) take data from a single very large recombining region such as an entire chromosome. In essence, we are currently only able to utilize information about the coalescent histories of the region in question—and this collection of histories may not match that of the entire population, which would be more accurately reflected in genome-wide data. In the Discussion, we address prospects for incorporating genome-scale data in demographic inference.

**Fig. 7:**
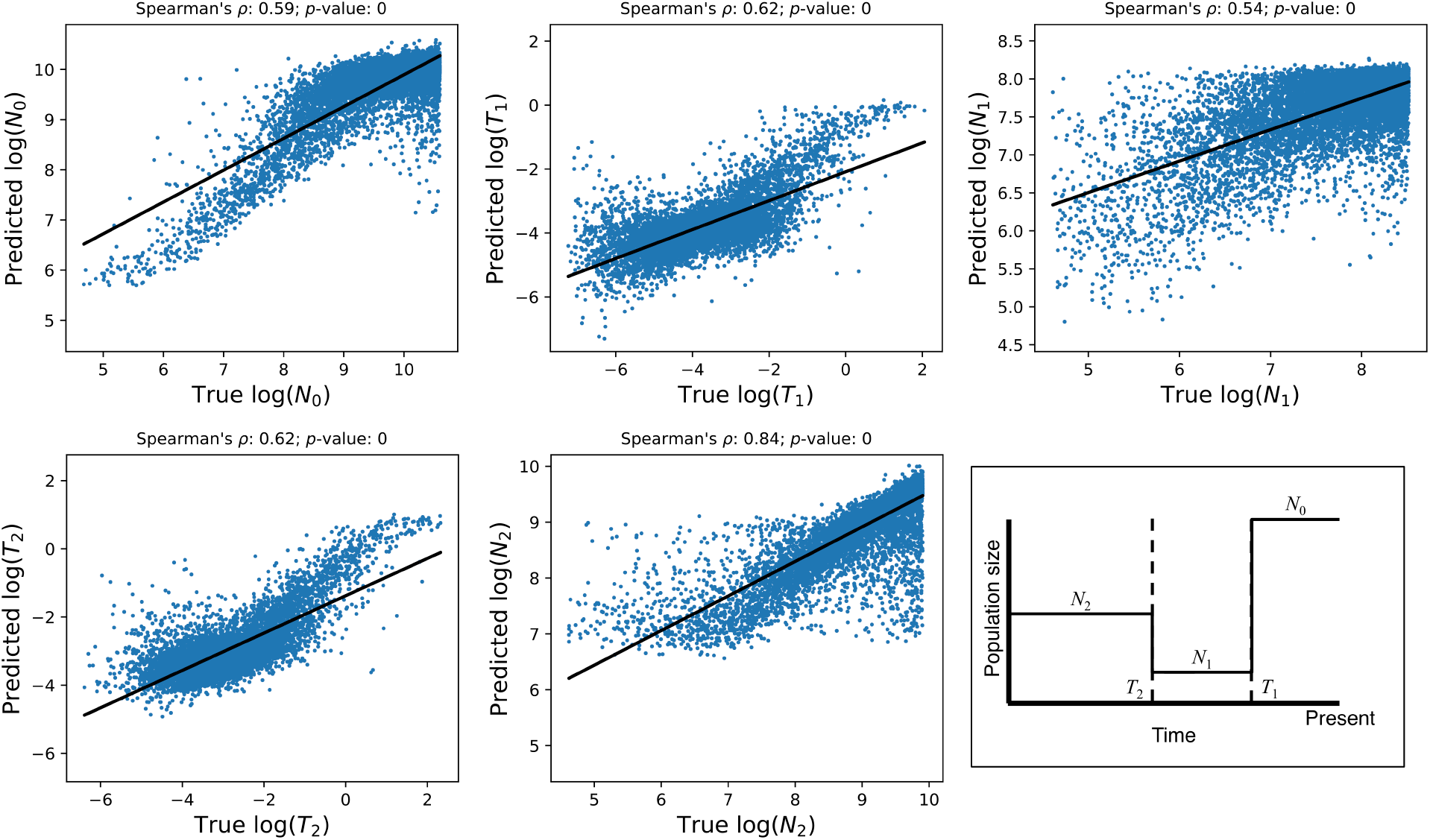
Accuracy of demographic inference CNN. The scatterplots show the correlation between true and predicted demographic parameter values using our best-performing CNN for this task when applied to an independent test set. Note that there may be some monotonicity in the relationship between the true and predicted values of some of these parameters, which may affect calculations of the Spearman correlation coefficients shown above each scatterplot. These estimates should thus be viewed as a rough summary of this relationship, while the RMSE values reported in the text better summarize our accuracy. The inset on the bottom right shows the demographic model and its five parameters.

## DISCUSSION

### Convolutional neural networks are well suited for population genetic problems

Population geneticists have devised a wide array of computational methods to make evolutionary inferences from genomic data. Typically the goal of these efforts is to aggregate information across genomic sites in order to make an accurate inference. These methods include likelihood-based approaches (e.g. Kim and Stephan 2002; Nielsen *et al.* 2005; Gutenkunst *et al.* 2009; Liu and Fu 2015), probabilistic graphical models such as hidden Markov models (e.g. Turner *et al.* 2005; Boitard *et al.* 2009; Lawson *et al.* 2012), and those that rely on the use one or more summary statistics designed to characterize patterns of variation within a genomic region (e.g. Tajima 1989; Fu and Li 1993; Kelly 1997; Fay and Wu 2000; Kim and Nielsen 2004; Voight *et al.* 2006; Ferrer-Admetlla *et al.* 2014). While these approaches differ substantially from one another, they all have one thing in common: they make use of population genomic theory to connect the features of a data set to the underlying evolutionary process. Here we have demonstrated the potential of an alternative approach: treating population genetic inference as an image recognition problem where the “image” is the population genetic alignment, which is directly fed as input to a CNN. In contrast to most mainstream approaches, this CNN approach makes use of the entirety of the data, rather than using theoretically derived estimators or closed-form likelihood functions to connect a small number of features of the data to an evolutionary process.

Here we have shown that CNNs perform remarkably well on a number of problems in population genetics. We developed CNNs with comparable if not greater power to detect selective sweeps, identify introgressed loci, and infer local recombination rates when compared to current methods on simulated data sets. The CNNs for detecting sweeps and introgression demonstrate the ability to use an alignment image to distinguish among multiple evolutionary models, while the recombination rate estimator demonstrates that continuous parameters can also be inferred. Finally, although our demographic parameter estimates were fairly imprecise, they were only based on a short stretch of the genome, and nonetheless demonstrate that CNNs have the potential to infer multiple parameters from a sequence alignment. While we were in the process of preparing this manuscript, Chan et al. completed an important study demonstrating that a CNN can accurately detect recombination hotspots (Chan *et al.* 2018). Taken together these results suggest that CNNs have enormous potential as a general paradigm for population genetic inference.

The effectiveness and generality of CNNs in population genetic inference should not be surprising. CNNs offer a number of intrinsic advantages that make them particularly amenable to population genetic data. First, there have been a number of efforts to move in the direction of making inferences on the basis of the full complement of data present in an alignment rather than one or more summary statistics (Li and Stephens 2003; Lawson *et al.* 2012; Smith *et al.* 2018). CNNs represent a natural way of examining the entirety of an alignment in order to increase inferential power. The development of novel CNN architectures to better handle spatial associations in the data across multiple scales (Yu and Koltun 2015) has the potential to improve CNN-driven population genetic inference even further. For example, improved ability to detect both the localized reduction in diversity at a sweep (Maynard Smith and Haigh 1974) as well as the potentially confounding skews in patterns of diversity produced in its flanking regions (Schrider *et al.* 2015) would be beneficial in sweep detection.

Another desirable property of CNNs is that they effectively perform automated feature detection (LeCun *et al.* 2015). Because they discover discriminatory information directly from the image, there is no need to manually construct an optimal set of features. CNNs may thus outperform methods based on a set of manually curated features as observed here, although this may not be the case for all tasks (e.g. Bellot *et al.* 2018). This brings up perhaps the strongest quality of CNNs in the context of evolutionary inference: because CNNs can make inference in the absence of statistics or a likelihood function, they can make predictions for phenomena for which there exists no analytical expectation.

Indeed, CNNs can tackle problems for which no relevant summary statistics have been devised—vectors of such statistics are required for other likelihood-free methods such as ABC (Beaumont 2010) or traditional supervised machine learning techniques (Schrider and Kern 2018). On a related note, neural networks are particularly amenable to the incorporation of disparate data types with no prior knowledge of their relationships. For example, here we have included both genotype information and positional information for segregating sites as branches to our networks, allowing both to be used together in prediction despite the fact that our network isn’t instructed how these two pieces of information relate to one another. All that is required is appropriate training data. Thus, we may not have to wait for theoretical advances in order to draw inferences from data, provided we are concerned with evolutionary models for which training data can be obtained from simulation—including the wide range of scenarios that could potentially be investigated via increasingly flexible and efficient forward simulators (Thornton 2014; Haller and Messer 2017; Kelleher *et al.* 2018).

This point is driven home by the success of our CNN for estimating recombination rates in autotetraploids from read pileup information alone—despite the input’s lack of genotype calls, let alone phased haplotypes, these inferences are nearly as accurate as those that we obtained from haplotype alignments. This result also suggests that CNNs may be well suited for other inferences where genotype calls are unreliable (e.g. low coverage sequencing data; Korneliussen *et al.* 2014) or unobtainable (e.g. pooled population sequencing; Schlötterer *et al.* 2014). Given CNNs’ flexibility, future studies should evaluate their potential to tackle not only those problems examined in this paper, but the myriad additional important challenges in evolutionary genetics to which they could be readily applied, including but not limited to uncovering adaptive introgression (Racimo *et al.* 2016), joint inference of selective and demographic histories (Sheehan and Song 2016), and even inferring structured outputs such as ancestral recombination graphs (Rasmussen *et al.* 2014).

### To what extent are CNNs robust to model misspecification?

Another particularly encouraging result of our recombination rate estimation analysis is that we were able to infer rates for data generated from a range of parameter values to which the CNN had not been exposed during training with very little decrease in accuracy. This ability to interpolate between training values is a particularly desirable property. First, it implies that CNNs can be used to create flexible inference tools using a modest training data set, and second that researchers can focus training between reasonable parameter bounds, without knowing the true (and often unknowable) underlying parameters; future efforts must explore the possibility of training networks to be robust to more extreme cases of model misspecification.

One illustrative example of the potential pitfalls of model misspecification is the problem of detecting selective sweeps without accounting for confounding demographic events. For example, population bottlenecks will skew genealogies in a manner similar to sweeps (Simonsen *et al.* 1995), and thus may result in a large fraction of false positives (Jensen *et al.* 2005; Nielsen *et al.* 2005). Schrider and Kern (2016) were able to mitigate this problem by designing a feature vector that is sensitive to the spatial skews in patterns of variation created by a sweep but insensitive to genome-wide skews produced by demographic events. Although this strategy is not possible with CNNs because they perform automated feature extraction, it may be that incorporating training examples generated under potentially confounding scenarios could alleviate this issue.

Therefore, future work must thoroughly 1) assess how CNNs trained on data simulated under one range of evolutionary parameters fare when applied to different parameterizations, and 2) determine whether robustness to such misspecification might be achieved by training a CNN under a wide range of parameter values that are likely to encapsulate the correct values—the recombination rate estimator’s successful interpolation suggests that this may be a possibility. Model misspecification is not a concern for tasks where training data may be obtained without simulation (e.g. detecting selective constraint; Schrider and Kern 2015), though in such cases one must take care to prevent dependencies between training and test examples because of shared evolutionary histories due to physical linkage or paralogy/orthology relationships (Washburn *et al.* 2018).

### Outstanding practical challenges associated with the application of CNNs to sequence data

Although the CNN approach outlined above has great potential, there are several outstanding challenges with applying CNNs to a wider spectrum of problems. One important obstacle is the large amount of training data required by CNNs, which makes applications requiring alignments of large regions (e.g. entire chromosomes) more difficult. This challenge includes both the generation of large labeled training examples, and time- and memory-efficient training with these large examples given limited computational resources. Fortunately, continued improvements in simulation speed (Kelleher *et al.* 2016; Kelleher *et al.* 2018) and the efficiency of CNN training (Chilimbi *et al.* 2014; Yu and Koltun 2015; Jouppi *et al.* 2017; Köster *et al.* 2017) is mitigating this problem. Such advances would be a boon for efforts to infer demographic parameters, which require simultaneously examining data sampled from across the genome or along an entire chromosome, unlike scans to infer locus-by-locus histories of selection/recombination/introgression. Advances in handling large or high-resolution images may also prove fruitful. For example, CNN-based strategies that simultaneously examine a number of smaller “patches”, each covering a portion of the image rather than the entirety of the image (e.g. Lu *et al.* 2015), may aid efforts to extract demographic information from genome-scale data.

Another challenge with the application of CNNs is that their performance can be sensitive to network architecture (Szegedy *et al.* 2015). There is no underlying theory for selecting optimal network architecture, though improved architectures are sure to continue to arise, and automated methods exist for optimizing the many hyperparameters of a given architecture (e.g. Snoek *et al.* 2012). Though we uncover some promising CNN architectures for population genetic inference, we suspect that substantial improvements can still be made.

We have also demonstrated that CNNs are sensitive to the input format of the population genetic alignment, and our work has yielded several insights along this front. First, we found that the ordering of haplotypes within the alignment can impact accuracy, and our results suggest that it is often beneficial to reorder haplotypes so that more similar chromosomes appear next to one another. This may be a suboptimal solution, and more creative approaches may be required to provide a more general strategy. To this end, research into permutation-invariant neural networks (Zaheer *et al.* 2017) may prove promising when dealing with sequence alignments. This is evidenced by Chan et al.’s recent findings that a permutation-invariant architecture improves both training speed and final accuracy of their CNN for detecting recombination rate hotspots (Chan *et al.* 2018). Chan et al.’s network avoids any convolution or pooling operations that combine information across individuals until an operation that collapses each column of the (filtered) alignment matrix down to a single value in an order-invariant manner (e.g. site-wise maximum). This design choice means that permuting the order of individuals within the alignment will have no impact on their network’s output. We also observed that 1D convolutions in the proper orientation perform as well as the more widely used 2D convolutions in many cases. Also, scaling response variables for regression problems (both log-scaling and standardization) may also affect accuracy. We therefore recommend that users experiment with these different ways of representing their data, as well as different CNN architectures, in order to find the design that works best for the task at hand.

Another important consideration of CNNs is that once trained, they are specialized to a particular problem as defined by the training set. That is, a CNN trained to infer recombination rates under a European demographic history may have reduced accuracy when applied to an African sample. Training under a variety of demographic scenarios may make a CNN more robust to this problem, but a question for further study is whether this can be accomplished without a loss in power relative to a more specialized CNN. Even a change as subtle as adding another chromosome to a dataset will make one of our previously trained CNNs inapplicable, as the input matrix would no longer be the proper size and either a new CNN must be trained or the data subsampled. Importantly, Chan et al. (2018) describe an architecture that can allow for variation in the number individuals in the input matrix. It is also important to note that in spite of their limitations, recent advances have greatly simplified training CNNs, and it will often be practical—or even preferable—for a researcher to create a CNN tailored to their specific data set.

### Are CNNs a black box?

Artificial neural networks are algorithms that seek to maximize their predictive accuracy by optimizing their internal mathematical operations on training data and CNNs are an extremely flexible subclass of these methods because they can act directly on the input data matrix (in our case a sequence alignment). However, one consequence of this is that CNNs are in some ways a “black box”. For example, a CNN cannot “explain” why it made a particular prediction given its input. Supervised machine learning algorithms in general have perhaps been unfairly maligned with this “black box” label. These methods can in principle reveal much about underlying processes by determining which features are most informative under certain scenarios (i.e. feature ranking; see Breiman 2001). For example, the observation that certain features are highly informative for recent but not ancient introgression (Schrider *et al.* 2018) suggests some key differences between the genealogies produced under these two scenarios. Due to their complex inner workings, less progress has been made in breaking through the CNN “black box” as compared to more traditional supervised machine learning techniques. However, some successful explanatory tools are available for CNNs (Ribeiro *et al.* 2016), and there is ongoing research in this area. Moreover, because the CNN framework we adopt here works on images, it may be possible to translate future breakthroughs in CNN interpretation from other fields (e.g. image recognition) into population genetic inference. Thus a more optimistic view is that as CNNs and related methods become more interpretable, these likelihood-free image recognition approaches may help to reveal theoretical insights into evolutionary processes.

In the near-term, CNNs may remain useful only as a predictive tool, and we will continue to rely on theoretical advances to improve our understanding of population genetic processes. In spite of the shortcomings noted above, the highly encouraging results that we have laid out here suggest that CNNs are able to discover information about the underlying genealogies from alignment images and to use this information to more accurately elucidate the evolutionary phenomena that have shaped these genealogies. CNNs have enormous potential for population genomic inference. We believe that progress on a host of problems could accelerate appreciably were this technology to be embraced by the field. Indeed, when it comes to the business-end of population genetics—drawing accurate evolutionary inferences from data—we predict that increasingly, likelihood-free approaches such as the ones we have describe here will prove most effective at solving existing problems, and expand the universe of problems that researchers can investigate.

## MATERIALS AND METHODS

### Computational environment for training CNNs

All CNNs used in this study were developed using two open source Python packages: Keras (version 2.0.6; https://keras.io/) to define neural network architecture and orchestrate training and testing, and TensorFlow (version 1.1.0; https://www.tensorflow.org/) as the backend (i.e. TensorFlow performs the computation during training/testing). CNN training is computationally intensive, but cloud-based GPU resources have made it affordable. As an example, our network for detecting selective sweeps was trained on a cloud-based system with one Nvidia K80 GPU. It took 6.6 hrs to train, and at $0.90 US dollars per hour the total cost was under $7. All code used for training is available online (https://github.com/flag0010/pop_gen_cnn).

### CNN validation strategy

For each task, we divided our simulated inputs into three sets: a training set, a validation set, and a test set. The training set was used to optimize the weights and biases of the CNN. The validation set was used during training to determine how well the CNN generalizes to unseen data, and adjustments were made to the CNN to improve its performance on the validation data. We also used the validation set to terminate training once accuracy on this set appeared to plateau—this process took different numbers of iterations for different tasks. Finally, the test set was used to obtain a performance assessment of the final trained network. Importantly, this test set was previously unseen by the CNN and therefore yields an unbiased evaluation of its accuracy. We used binom.test in R to estimate 95% confidence intervals for classification accuracies.

### Evaluating techniques for rescaling and reordering inputs to improve CNN accuracy

To evaluate the impact of alternative data preparation techniques, we developed a simple CNN that estimates the locus-wide population mutation rate *θ*=4*NμL* where *μ* is the mutation rate per base pair per generation and *L* is the physical length of the locus being examined. This CNN is trained using alignment images with forty chromosomes and *θ* drawn uniformly between 10 and 50 as simulated for a panmictic, constant sized population by ms (Hudson 2002). We trained this CNN to minimize the root mean squared error (RMSE) between its prediction and the true value of *θ* using 4,000 training matrices. Then its accuracy was scored on 1,000 test matrices that the CNN was never trained on. These values were compared under different data preparation approaches described below.

First, the matrices output by most coalescent simulation software, including ms, encode ancestral and derived alleles for bialleleic sites as 0 and 1, respectively, and present the matrix with phased haploid chromosomes as rows and sites as columns. When doing 1D convolutions, we sought to use row-wise convolutional filters (Fig. 1C), i.e. those that examine each chromosome in our sample across a small number of contiguous segregating sites (specified by the “kernel_size” parameter in Keras) before sliding the filter forward one site (our stride length, “strides” in Keras, was always set to 1). At present Keras does not allow for row-wise 1D convolutions, so we accomplished this by transposing the alignment matrix and performing column-wise convolutions.

We also assessed the impact on accuracy of sorting the chromosomes in the alignment by genetic similarity. For example, the matrices in Fig. 2 contain identical information, but chromosomes in the matrix on the left are randomized, while on the right they are sorted by genetic similarity. We offer a fast algorithm for sorting matrices by genetic similarity (https://github.com/flag0010/pop_gen_cnn/blob/master/sort.min.diff.py).

### Introgression detection

To detect introgression, we simulated phased haploid training and test examples with msmove (https://github.com/geneva/msmove) from the same demographic model that Schrider et al. (2018) used to train the FILET classifier for detecting introgression between *Drosophila simulans* and *D. sechellia*. In total we produced 237,500 coalescent simulations from 3 classes: 112,500 without no migration between species (No Introgression), 112,500 with gene flow from *D. simulans* into *D. sechellia* (*sim*→*sech*), and 12,500 with gene flow from *D. sechellia* into *D. simulans* (*sech*→*sim*). We used fewer *sech*→*sim* examples because test runs on smaller training sets suggested that the network could detect this class fairly accurately, which allowed us to increase the sampling of the other two more challenging classes by simulating more examples from them. To our knowledge this approach of intentionally inflating the number and proportion of training examples from the more challenging classes is unusual, as typically a balanced training set is preferred. However we found that including additional examples from classes into our data set substantially improved our ability to correctly them. The simulations were randomly assigned to training and validation sets so that the training set included 107,500 examples each from the No Introgression and *sim*→*sech* classes, and 7,500 examples from the *sech*→*sim* class. Both the validation set and the test set contained 2,500 of each class (i.e. 7,500 total). Importantly, because our test and validation sets were evenly balanced, they provided unbiased estimates of our accuracy.

As in the *Drosophila* data set to which Schrider et al. applied FILET, each of our coalescent simulations generated 34 chromosomes (14 *D. sechellia* and 20 *D. simulans*). Each column in the alignment corresponded to a biallelic polymorphism, which was encoded as “0” (ancestral allele) or “1” (derived allele) for each chromosome. In practice, the ancestral and derived states may not be known with 100% certainty, and one may instead use major/minor alleles, or randomly mispolarize a fraction of sites in the training data if one has an estimate of the fraction of mispolarized sites in the true data. The effects of these design choices on performance may then be evaluated on test data. Each matrix was organized so that individual chromosomes were grouped by species. Each coalescent simulation produced a different number of segregating sites (with the largest containing 1201 polymorphisms). Because the CNN’s input matrices must all have the same dimensions, we padded the right side of all matrices with fewer than 1201 polymorphisms with columns containing only “0” until the total number of columns reached 1201. Finally we transposed this matrix resulting in a 1201×34 matrix for each coalescent simulation. In practice, one will have to set the image width to the largest number of SNPs encountered across all training, test/validation, or real data examples included in the analysis. Alternatively, one may select a fixed number of segregating sites to include in the analysis, in which case each example may correspond to a different physical size (creating additional variance in total recombination rates). Thus, when using this alternative approach, one should adjust the lengths of simulated examples accordingly.

We trained a CNN architecture with three 1D-convolutional layers (kernel size = 2), each followed by average-pooling, and finally two densely connected layers (i.e. the same network architecture as the main network branch illustrated in Fig. 1C, but with one additional dense layer). These layers contained 256, 128, 128, 128, and 128 neurons, respectively. To avoid overfitting during training, each layer used dropout regularization (randomly removing 25% of neurons between convolutional layers during each training iteration, and 50% between densely connected layers) and rectified linear unit activation functions (i.e. ReLUs; Hahnloser *et al.* 2000; Nair and Hinton 2010). Dropout regularization encourages the CNN to learn redundant representations of the data, thereby reducing the network’s dependence on individual weights (Srivastava *et al.* 2014). The last layer was a sigmoid output layer with 3 neurons, each corresponding to the 3 classes given above. The CNN was trained using the Adam optimization procedure (Kingma and Ba 2014), a categorical cross-entropy loss function, and a mini-batch size of 256. The CNN was run for 19 training iterations through the training data.

### Recombination rate: phased haplotype version

For the recombination rate estimator we used ms (Hudson 2002) to simulate 50 phased chromosomes, each with a target length of 20kb. To do so, we drew a population size (*N*) from the following values: 5,000, 10,000, 15,000, 20,000, and 50,000, and set the population-scaled mutation rate parameter 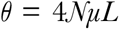 (letting *μ*=1.5×10^-8^ and *L*=20kb). We also set a population-scaled recombination rate, 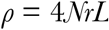, where *r* is the per bp crossover rate per meiosis, by drawing *r* from a bounded exponential distribution raging from10^-8^ to 10^-6^. This yields a range of *ρ* per base pair of 2×10^-4^ to 2×10^-1^. These values roughly encompass the range of recombination rates experienced in humans and *Drosophila*. Following this procedure, we generated 156,275 coalescent simulations. ~92% were used to train the CNN, and ~4% each were set aside for validation and testing. To assess our CNNs ability to interpolate to unseen population sizes, we also created 5,000 additional test matrices using the procedures above, but with 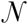 drawn uniformly from the following: 30,000, 35,000, 40,000, and 45,000.

Each simulation was represented by a matrix of 50 rows, one for each chromosome, and 418 columns (the largest number of segregating sites). As before, we encoded the ancestral allele with “0” and the derived allele with “1”. Because not all simulations resulted in the same number of polymorphisms, we padded both the genotype matrix and the position vector in the same manner as for the introgression CNN, bringing the total size of each matrix to 50×418. Next, we sorted each matrix by genetic similarity among chromosomes as described above and then transposed the matrix to 418×50. We also extracted the segregating site positions vector from the ms output which represents each position as a real number between zero (the leftmost position on the simulated chromosome) and one (the rightmost position). For simulations with fewer than 418 segregating sites, we padded the positions vector with “-1”s.

We transformed the *ρ* values for the training, validation, and test sets by taking the natural log of each value and centering them on the mean of the training set. By using the mean from the training set for all transformations, we ensure that there is no leakage of information between training and validation/testing.

We trained a CNN with two input branches. The first branch took the haplotype matrices as input and included three 1D-convolutional layers (kernel size = 2), each followed by average-pooling. These layers contained 1250, 256, and 256 neurons, respectively. Each of these layers uses dropout normalization (25%), L2-regularization of the weights (λ = 0.0001), and ReLU activation functions. The second branch took the position vector as input and contains one densely connected layer with 64 neurons, again using dropout normalization (10%) and a ReLU activation function. The two branches are then merged into another densely connected layer of 256 neurons with ReLU activation functions. Finally, the output layer is a single neuron with a simple linear activation function that predicts the continuous *ρ* value. The CNN was trained using the Adam optimization algorithm, using mean-squared error as our loss function, and a mini-batch size of 32. The CNN was trained for 16 iterations.

We compared our CNN’s results to those of LDhat version 2.2a (https://github.com/auton1/LDhat). We chose LDhat because it is widely used to estimate historical recombination rates, and because it can be efficiently run on large data sets. LDhat will estimate *ρ* only for a specified population mutation rate 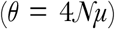, and we supplied it with the exact *θ* value used for each coalescent simulation. This was done by creating five likelihood lookup tables using the complete program, all set for 50 haploid chromosomes, for the following *θ* values: 6, 12, 18, 24, and 60. Respectively, these correspond to 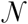 = 5,000, 10,000, 15,000, 20,000, and 50,000 (the same values we used for training our CNNs). LDhat only predicts values within the bounds of the lookup table. Therefore, to facilitate a fair comparison to results from our CNN, which is unbounded, we selected the maximum *ρ* value in the likelihood lookup table to be 133.3% of the true maximum for each *θ*. We then set the grid size of *ρ* equal 1, and estimated *ρ* on the test set using LDhat’s pairwise program.

In contrast, the CNN was not provided information about *θ*, and instead had to infer *ρ* independent of *θ*. This ability would be a desirable property for an estimator, as *θ* is likely to vary considerably across the genome and outside of simulated data sets one may never know *θ* precisely. On the other hand, the CNN was provided with the physical distance between segregating sites, information LDhat does not utilize but which will generally be available when making inferences on real data. Both of these factors make our direct comparison of the CNN with LDhat imperfect because each had access to information the other lacked when producing its estimate. Nonetheless we consider this example a useful illustration of the CNN’s performance.

### Recombination rate: autotetraploid version

We sought to train a CNN to estimate a locus-wide recombination rate in autotetraploid genomes. To add a level of methodological realism to this problem, we did so from a matrix storing a simple summary of read pileup information at each site for each individual.

To this end, we generated new coalescent simulations with 48 chromosomes each following the procedure outlined above for the haploid CNN. This approach is reasonable because it has been shown that the standard coalescent approximates the appropriate coalescent for autotetraploids as long as *N* is larger than a few hundred (Arnold *et al.* 2012). We generated 217,500 coalescent simulations, and randomly assigned 200,000 to the training set, 10,000 to the validation set, and 7,500 to the test set. Next, within each coalescent simulation, we randomly partitioned our 48 chromosomes into twelve sets of four. Each set represents one synthetic autotetraploid genome and every site has five possible genotypes (*AAAA*, *AAAa*, *AAaa*, *Aaaa*, and *aaaa*). For each autotetraploid genome *i* and each site *j* we simulated the number of reads covering the site (*C*_*ij*_) by drawing a random sample from a Poisson distribution with λ = 25. Then we selected the number of reads representing the *a* allele *R_ij_ ~ Binom(n=C_ij_, p=x_ij_)*, where *x*_*ij*_ represents the frequency of the *a* allele in the tetraploid genotype (i.e. 0, 0.25, 0.5, 0.75, and 1 for the five genotypes listed above). For each individual *i* at site *j*, the corresponding entry in the input matrix was the fraction *R*_*ij*_/*C*_*ij*_, i.e. the fraction of reads supporting the derived allele. The *AAAA* and *aaaa* genotypes were always 0 and 1, respectively. For the three heterozygous genotypes (*AAAa*, *AAaa*, and *Aaaa*), *R*_*ij*_/*C*_*ij*_ varied based on sampling error but had expected values of 0.25, 0.5, and 0.75, respectively. Thus at each site the original 48 chromosomes were reduced to a set of 12 values corresponding to the fractions of reads supporting the *a* allele in a pool of sequence reads from an autotetraploid sequenced at ~25X coverage. Note that this scheme includes neither sequencing error, nor the site-specific depth which would be necessary to calculate a likelihood, but is nonetheless adequate for our proof of concept.

As above, we sorted the rows of this matrix by genetic similarity and padded each matrix with zeros to a length of 460 (the most segregating sites of any of the simulated matrices) before transposing, yielding a 460×12 matrix. Again, we recorded the padded vector of positions from the simulation output. Our CNN architecture was identical to the one given above for the phased haplotype version, except for the dimensionality of the input changed to 460×12, and we reduced the first convolutional layer from 1250 to 256 because of the smaller second dimension of the input. The CNN was trained for 9 iterations.

### Detecting selective sweeps and discriminating between modes of selection

For detecting selective sweeps, we used the same coalescent simulations that Schrider and Kern (2017) used to train a classifier to detect sweeps in the JPT population (Japanese individuals from Tokyo) from Phase 3 of the 1000 Genomes dataset (Auton *et al.* 2015). The JPT demographic scenario is one where detecting selective sweeps is fairly difficult (see Figure S1 from Schrider and Kern 2017), as expected for bottlenecked populations (Jensen *et al.* 2005). For this CNN, we began with a set of 269,000 simulated genomic windows with the 5 following classes: a recent hard sweep (i.e. fixation of a *de novo* beneficial mutation), a recent soft sweep (i.e. fixation of a beneficial but previously neutral segregating polymorphism), a region linked to a nearby hard sweep, a region linked to a nearby soft sweep, and a neutrally evolving region. Each simulated alignment contained 208 chromosomes and we kept only coalescent simulations that contained ≤ 5,000 segregating sites, and again padded with zeros so that all matrices were 208×5000. This left 238,655 simulations, and from those we constructed a training set of 233,655 simulations. In trial runs, we found that regions flanking hard and soft sweeps were the most difficult classes to predict, so we again simulated additional examples from these more challenging classes. This shifted the balance of our training set so that is was comprised of approximately 13% neutral regions, 17% each for hard and soft sweeps, and 26.5% each for regions linked to nearby hard and soft sweeps windows. We then set aside an evenly balanced set of 2,000 simulations for validation and 3,000 for testing.

As before, we sorted each matrix by genetic similarity among chromosomes and then transposed the matrix to 5000×208. We also extracted the segregating site positions vector from these simulations which were generated by discoal (Kern and Schrider 2016), which like ms represents each position as a real number between zero and one.

As above, we trained a CNN with two input branches. The first branch took the haplotype matrices as input and included five 1D-convolutional layers (kernel size = 2), each followed by average-pooling. These layers each contained 256 neurons and used dropout normalization (20%). The second branch took the position vector as input and contained one densely connected layer with 64 neurons, again using dropout normalization (10%). The two branches were then merged into another densely connected layer of 256 neurons with 25% dropout. Each hidden layer of the network used L2-regularization of the weights (λ = 0.0001) and ReLU as the activation function. Finally, the output of this layer was fed to a five neuron layer with softmax activation functions that predicts the five classes given above. The CNN was trained using the Adam optimization algorithm, the categorical cross-entropy loss function, and a mini-batch size of 32. The CNN was trained for 3 iterations.

### Inferring population size histories

To show how CNNs can be used to infer species’ demographic histories, and how CNN architecture can impact this inference, we experimented with a variety of CNN approaches to infer the 5 parameters of a 3-epoch model of instantaneous population size changes (i.e. 3 population sizes and 2 times of size change). We also use this challenging problem as an opportunity to evaluate how alternative approaches to building a CNN can influence its performance. In effect, we conducted a full grid search of the following attributes of both our CNN architecture and input/output format: the dimensionality of our convolutions (1D or 2D), the kernel size (i.e. the width of our 1D convolutional filters and both the height and width of our square 2D filters; we tried each multiple of 2 raging from 2 to 10), whether to include dropout (yes or no) following max pooling steps or dense layers, whether to sort our rows based on similarity (yes or no), whether to log-scale our response variables (yes or no), and whether to represent ancestral and derived alleles as -1/1 or as 0/255. When included, our dropout layers immediately followed both max pooling steps, the dense layer following the distance input layer, and the final dense layer. Each of these dropout steps randomly removed 25% of neurons. Each response variable was transformed to a 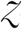-score according to the sample mean and variance for that variable across all simulated examples.

The network we used for this task had two branches: a standard CNN like that depicted in Fig. 1B–C but with more convolutional layers (four CNN layers each producing 128 filters and each followed by a max pooling layer with a kernel size of 2), and a dense neural network layer (consisting of 32 nodes) taking positional information as its input, and concatenating its output with that of the final max pooling layer of the CNN prior to being fed into the final dense layer (256 nodes). The positional information was a vector, ***d***, whose length was the maximum of the number of segregating sites observed across all simulated examples minus one. Each value in the vector *d*_*i*_ was simply the distance (scaled between zero and one where one is the total length of the simulated region) between segregating site *i* and site *i*-1.

In total, we simulated 100,000 alignments of phased chromosomes using ms. 10,000 each were set aside for testing and validation, while the remaining 80,000 were used for training. The simulated population size histories were generated randomly—each demographic model parameter was drawn uniformly from a range listed in supplementary table S2. Each simulated region was roughly equivalent 1.5 Mbp in the human genome, assuming per base pair mutation and recombination rates of 1.2×10^-8^ and 1×10^-8^, respectively. However, in order to make the size of the simulation output more tractable for processing in a CNN we divided the mutation rate by 10 (equivalent to randomly downsampling the number of polymorphisms included in the input by a factor of 10). During training we used a batch size of 200, trained our networks for up to 10 iterations, and retained the best performing CNN as assessed on the validation set. Often the best CNN was obtained prior to completing all 10 training iterations. We then evaluated the performance of the best CNN for each network architecture and input format on the test set by calculating total RMSE (our loss function for this task); we also calculated Spearman correlation coefficients between the true and predicted values for each of the five demographic model parameters.

## Supporting information

## ACKNOWLEDGMENTS

We thank Matt Hahn for comments on the manuscript. DRS was funded by the National Institutes of Health under award number R00HG008696. YB would like to acknowledge the Minnesota Supercomputing Institute for computational resources.

## SUPPLEMENTARY TABLE LEGENDS

**Supplementary table S1: The effect of different neural network input/output/architecture hyperparameters on demographic inference error.**

**Supplementary table S2: Demographic parameter ranges used to simulate 3-epoch population size histories.**

## REFERENCES

Arnold, B., K. Bomblies and J. Wakeley, 2012 Extending coalescent theory to autotetraploids. Genetics 192: 195–204.

Auton, A., L. D. Brooks, R. M. Durbin, E. P. Garrison, H. M. Kang et al., 2015 A global reference for human genetic variation. Nature 526: 68–74.

Auton, A., and G. McVean, 2007 Recombination rate estimation in the presence of hotspots. Genome Res. 17: 1219–1227.

Beaumont, M. A., 2010 Approximate Bayesian computation in evolution and ecology. Annual review of ecology, evolution, and systematics 41: 379–406.

Begun, D. J., and C. F. Aquadro, 1992 Levels of naturally occurring DNA polymorphism correlate with recombination rates in *D. melanogaster*. Nature 356: 519–520.

Begun, D. J., A. K. Holloway, K. Stevens, L. W. Hillier, Y.-P. Poh et al., 2007 Population genomics: whole-genome analysis of polymorphism and divergence in Drosophila simulans. PLoS Biol. 5: e310.

Bellot, P., G. de los Campos and M. Pérez-Enciso, 2018 Can Deep Learning Improve Genomic Prediction of Complex Human Traits? Genetics: genetics. 301298.302018.

Boitard, S., C. Schlötterer and A. Futschik, 2009 Detecting selective sweeps: a new approach based on hidden Markov models. Genetics 181: 1567–1578.

Brandvain, Y., A. M. Kenney, L. Flagel, G. Coop and A. L. Sweigart, 2014 Speciation and introgression between Mimulus nasutus and Mimulus guttatus. PLoS Genet. 10: e1004410.

Breiman, L., 2001 Statistical modeling: The two cultures (with comments and a rejoinder by the author). Statistical science 16: 199–231.

Chan, A. H., P. A. Jenkins and Y. S. Song, 2012 Genome-wide fine-scale recombination rate variation in Drosophila melanogaster. PLoS Genet. 8: e1003090.

Chan, J., V. Perrone, J. P. Spence, P. A. Jenkins, S. Mathieson et al., 2018 A Likelihood-Free Inference Framework for Population Genetic Data using Exchangeable Neural Networks. bioRxiv.

Charlesworth, B., M. Morgan and D. Charlesworth, 1993 The effect of deleterious mutations on neutral molecular variation. Genetics 134: 1289–1303.

Chilimbi, T. M., Y. Suzue, J. Apacible and K. Kalyanaraman, 2014 Project Adam: Building an Efficient and Scalable Deep Learning Training System, pp. 571–582 in OSDI.

Corbett-Detig, R., and R. Nielsen, 2017 A hidden Markov model approach for simultaneously estimating local ancestry and admixture time using next generation sequence data in samples of arbitrary ploidy. PLoS Genet. 13: e1006529.

Dieleman, S., and B. Schrauwen, 2014 End-to-end learning for music audio, pp. 6964–6968 in Acoustics, Speech and Signal Processing (ICASSP), 2014 IEEE International Conference on. IEEE.

Dutheil, J. Y., G. Ganapathy, A. Hobolth, T. Mailund, M. K. Uyenoyama et al., 2009 Ancestral population genomics: the coalescent hidden Markov model approach. Genetics 183: 259–274.

Elyashiv, E., S. Sattath, T. T. Hu, A. Strutsovsky, G. McVicker et al., 2016 A genomic map of the effects of linked selection in Drosophila. PLoS Genet. 12: e1006130.

Ewing, G., and J. Hermisson, 2010 MSMS: a coalescent simulation program including recombination, demographic structure and selection at a single locus. Bioinformatics 26: 2064–2065.

Fay, J. C., and C.-I. Wu, 2000 Hitchhiking under positive Darwinian selection. Genetics 155: 1405–1413.

Ferrer-Admetlla, A., M. Liang, T. Korneliussen and R. Nielsen, 2014 On detecting incomplete soft or hard selective sweeps using haplotype structure. Mol. Biol. Evol. 31: 1275–1291.

Fontaine, M. C., J. B. Pease, A. Steele, R. M. Waterhouse, D. E. Neafsey et al., 2015 Extensive introgression in a malaria vector species complex revealed by phylogenomics. Science 347: 1258524.

Fu, Y.-X., and W.-H. Li, 1993 Statistical tests of neutrality of mutations. Genetics 133: 693–709.

Gao, F., C. Ming, W. Hu and H. Li, 2016 New software for the fast estimation of population recombination rates (FastEPRR) in the genomic era. G3: Genes, Genomes, Genetics 6: 1563–1571.

Garrigan, D., S. B. Kingan, A. J. Geneva, P. Andolfatto, A. G. Clark et al., 2012 Genome sequencing reveals complex speciation in the Drosophila simulans clade. Genome Res. 22: 1499–1511.

Garud, N. R., P. W. Messer, E. O. Buzbas and D. A. Petrov, 2015 Recent selective sweeps in North American Drosophila melanogaster show signatures of soft sweeps. PLoS Genet. 11: e1005004.

Gazave, E., L. Ma, D. Chang, A. Coventry, F. Gao et al., 2014 Neutral genomic regions refine models of recent rapid human population growth. Proceedings of the National Academy of Sciences 111: 757–762.

Geneva, A. J., C. A. Muirhead, S. B. Kingan and D. Garrigan, 2015 A new method to scan genomes for introgression in a secondary contact model. PLoS ONE 10: e0118621.

Gutenkunst, R. N., R. D. Hernandez, S. H. Williamson and C. D. Bustamante, 2009 Inferring the joint demographic history of multiple populations from multidimensional SNP frequency data. PLoS Genet. 5: e1000695.

Hahn, M. W., 2018 Molecular Population Genetics. Oxford University Press.

Hahnloser, R. H., R. Sarpeshkar, M. A. Mahowald, R. J. Douglas and H. S. Seung, 2000 Digital selection and analogue amplification coexist in a cortex-inspired silicon circuit. Nature 405: 947.

Haller, B., and P. Messer, 2017 SLiM 2: Flexible, Interactive Forward Genetic Simulations. Mol. Biol. Evol. 34: 230.

Hedrick, P. W., 2013 Adaptive introgression in animals: examples and comparison to new mutation and standing variation as sources of adaptive variation. Mol. Ecol. 22: 4606–4618.

Hellenthal, G., G. B. Busby, G. Band, J. F. Wilson, C. Capelli et al., 2014 A genetic atlas of human admixture history. Science 343: 747–751.

Hermisson, J., and P. S. Pennings, 2005 Soft sweeps molecular population genetics of adaptation from standing genetic variation. Genetics 169: 2335–2352.

Hey, J., 2009 Isolation with migration models for more than two populations. Mol. Biol. Evol. 27: 905–920.

Hey, J., and J. Wakeley, 1997 A coalescent estimator of the population recombination rate. Genetics 145: 833–846.

Hill, W. G., and A. Robertson, 1966 The effect of linkage on limits to artificial selection. Genetics Research 8: 269–294.

Hobolth, A., O. F. Christensen, T. Mailund and M. H. Schierup, 2007 Genomic relationships and speciation times of human, chimpanzee, and gorilla inferred from a coalescent hidden Markov model. PLoS Genet. 3: e7.

Hornik, K., 1991 Approximation capabilities of multilayer feedforward networks. Neural networks 4: 251–257.

Hudson, R. R., 1987 Estimating the recombination parameter of a finite population model without selection. Genetics Research 50: 245–250.

Hudson, R. R., 2001 Two-locus sampling distributions and their application. Genetics 159: 1805–1817.

Hudson, R. R., 2002 Generating samples under a Wright–Fisher neutral model of genetic variation. Bioinformatics 18: 337–338.

Hudson, R. R., and N. L. Kaplan, 1985 Statistical properties of the number of recombination events in the history of a sample of DNA sequences. Genetics 111: 147–164.

Jensen, J. D., Y. Kim, V. B. DuMont, C. F. Aquadro and C. D. Bustamante, 2005 Distinguishing between selective sweeps and demography using DNA polymorphism data. Genetics 170: 1401–1410.

Joly, S., P. A. McLenachan and P. J. Lockhart, 2009 A statistical approach for distinguishing hybridization and incomplete lineage sorting. The American Naturalist 174: E54–E70.

Jouppi, N. P., C. Young, N. Patil, D. Patterson, G. Agrawal et al., 2017 In-datacenter performance analysis of a tensor processing unit, pp. 1–12 in Proceedings of the 44th Annual International Symposium on Computer Architecture. ACM.

Kaplan, N. L., R. Hudson and C. Langley, 1989 The “hitchhiking effect” revisited. Genetics 123: 887–899.

Kelleher, J., A. M. Etheridge and G. McVean, 2016 Efficient coalescent simulation and genealogical analysis for large sample sizes. PLoS Comput. Biol. 12: e1004842.

Kelleher, J., K. Thornton, J. Ashander and P. Ralph, 2018 Efficient pedigree recording for fast population genetics simulation. bioRxiv: 248500.

Kelly, J. K., 1997 A test of neutrality based on interlocus associations. Genetics 146: 1197–1206.

Kern, A. D., and D. Haussler, 2010 A population genetic hidden Markov model for detecting genomic regions under selection. Mol. Biol. Evol. 27: 1673–1685.

Kern, A. D., and D. R. Schrider, 2016 discoal: flexible coalescent simulations with selection. Bioinformatics 32: btw556.

Kim, Y., 2014 Convolutional neural networks for sentence classification. arXiv preprint arXiv:1408.5882.

Kim, Y., and R. Nielsen, 2004 Linkage disequilibrium as a signature of selective sweeps. Genetics 167: 1513–1524.

Kim, Y., and W. Stephan, 2002 Detecting a local signature of genetic hitchhiking along a recombining chromosome. Genetics 160: 765–777.

Kingma, D. P., and J. Ba, 2014 Adam: A method for stochastic optimization. arXiv preprint arXiv:1412.6980.

Kong, A., G. Thorleifsson, D. F. Gudbjartsson, G. Masson, A. Sigurdsson et al., 2010 Fine-scale recombination rate differences between sexes, populations and individuals. Nature 467: 1099–1103.

Korneliussen, T. S., A. Albrechtsen and R. Nielsen, 2014 ANGSD: analysis of next generation sequencing data. BMC Bioinformatics 15: 356.

Köster, U., T. Webb, X. Wang, M. Nassar, A. K. Bansal et al., 2017 Flexpoint: An adaptive numerical format for efficient training of deep neural networks, pp. 1742–1752 in Advances in Neural Information Processing Systems.

Krizhevsky, A., I. Sutskever and G. E. Hinton, 2012 Imagenet classification with deep convolutional neural networks, pp. 1097–1105 in Advances in neural information processing systems.

Kulathinal, R. J., L. S. Stevison and M. A. Noor, 2009 The genomics of speciation in Drosophila: diversity, divergence, and introgression estimated using low-coverage genome sequencing. PLoS Genet. 5: e1000550.

Langley, C. H., K. Stevens, C. Cardeno, Y. C. G. Lee, D. R. Schrider et al., 2012 Genomic variation in natural populations of *Drosophila melanogaster*. Genetics 192: 533–598.

Lawrence, S., C. L. Giles, A. C. Tsoi and A. D. Back, 1997 Face recognition: A convolutional neural-network approach. IEEE transactions on neural networks 8: 98–113.

Lawson, D. J., G. Hellenthal, S. Myers and D. Falush, 2012 Inference of population structure using dense haplotype data. PLoS Genet. 8: e1002453.

LeCun, Y., Y. Bengio and G. Hinton, 2015 Deep learning. Nature 521: 436–444.

LeCun, Y., L. Bottou, Y. Bengio and P. Haffner, 1998 Gradient-based learning applied to document recognition. Proceedings of the IEEE 86: 2278–2324.

Li, H., and R. Durbin, 2011 Inference of human population history from individual whole-genome sequences. Nature 475: 493–496.

Li, N., and M. Stephens, 2003 Modeling linkage disequilibrium and identifying recombination hotspots using single-nucleotide polymorphism data. Genetics 165: 2213–2233.

Lin, K., A. Futschik and H. Li, 2013 A fast estimate for the population recombination rate based on regression. Genetics 194: 473–484.

Lin, K., H. Li, C. Schlötterer and A. Futschik, 2011 Distinguishing positive selection from neutral evolution: boosting the performance of summary statistics. Genetics 187: 229–244.

Lipson, M., P.-R. Loh, A. Levin, D. Reich, N. Patterson et al., 2013 Efficient moment-based inference of admixture parameters and sources of gene flow. Mol. Biol. Evol. 30: 1788–1802.

Liu, X., and Y.-X. Fu, 2015 Exploring population size changes using SNP frequency spectra. Nat. Genet. 47: 555–559.

Loh, P.-R., M. Lipson, N. Patterson, P. Moorjani, J. K. Pickrell et al., 2013 Inferring admixture histories of human populations using linkage disequilibrium. Genetics 193: 1233–1254.

Lu, X., Z. Lin, X. Shen, R. Mech and J. Z. Wang, 2015 Deep multi-patch aggregation network for image style, aesthetics, and quality estimation, pp. 990–998 in Proceedings of the IEEE International Conference on Computer Vision.

Marth, G. T., E. Czabarka, J. Murvai and S. T. Sherry, 2004 The allele frequency spectrum in genome-wide human variation data reveals signals of differential demographic history in three large world populations. Genetics 166: 351–372.

Martin, S. H., K. K. Dasmahapatra, N. J. Nadeau, C. Salazar, J. R. Walters et al., 2013 Genome-wide evidence for speciation with gene flow in Heliconius butterflies. Genome Res. 23: 1817–1828.

Maynard Smith, J., and J. Haigh, 1974 The hitch-hiking effect of a favourable gene. Genet. Res. 23: 23–35.

McVean, G. A., S. R. Myers, S. Hunt, P. Deloukas, D. R. Bentley et al., 2004 The fine-scale structure of recombination rate variation in the human genome. Science 304: 581–584.

Mitchell, T. M., 1997 Artificial neural networks. Machine Learning 45: 81–127.

Nair, V., and G. E. Hinton, 2010 Rectified linear units improve restricted boltzmann machines, pp. 807–814 in Proceedings of the 27th international conference on machine learning (ICML-10).

Nielsen, R., and J. Wakeley, 2001 Distinguishing migration from isolation: a Markov chain Monte Carlo approach. Genetics 158: 885–896.

Nielsen, R., S. Williamson, Y. Kim, M. J. Hubisz, A. G. Clark et al., 2005 Genomic scans for selective sweeps using SNP data. Genome Res. 15: 1566–1575.

Pavlidis, P., J. D. Jensen and W. Stephan, 2010 Searching for footprints of positive selection in whole-genome SNP data from nonequilibrium populations. Genetics 185: 907–922.

Price, A. L., A. Tandon, N. Patterson, K. C. Barnes, N. Rafaels et al., 2009 Sensitive detection of chromosomal segments of distinct ancestry in admixed populations. PLoS Genet. 5: e1000519.

Pudlo, P., J.-M. Marin, A. Estoup, J.-M. Cornuet, M. Gautier et al., 2016 Reliable ABC model choice via random forests. Bioinformatics 32: 859–866.

Pybus, M., P. Luisi, G. M. Dall’Olio, M. Uzkudun, H. Laayouni et al., 2015 Hierarchical boosting: a machine-learning framework to detect and classify hard selective sweeps in human populations. Bioinformatics 31: 3946–3952.

Racimo, F., D. Marnetto and E. Huerta-Sanchez, 2016 Signatures of archaic adaptive introgression in present-day human populations. Mol. Biol. Evol. 34: 296–317.

Rasmussen, M. D., M. J. Hubisz, I. Gronau and A. Siepel, 2014 Genome-wide inference of ancestral recombination graphs.

Ribeiro, M. T., S. Singh and C. Guestrin, 2016 Why should i trust you?: Explaining the predictions of any classifier, pp. 1135–1144 in Proceedings of the 22nd ACM SIGKDD International Conference on Knowledge Discovery and Data Mining. ACM.

Ronen, R., N. Udpa, E. Halperin and V. Bafna, 2013 Learning natural selection from the site frequency spectrum. Genetics 195: 181–193.

Rosenzweig, B. K., J. B. Pease, N. J. Besansky and M. W. Hahn, 2016 Powerful methods for detecting introgressed regions from population genomic data. Mol. Ecol. 25: 2387–2397.

Rumelhart, D. E., G. E. Hinton and R. J. Williams, 1986 Learning representations by back-propagating errors. Nature 323: 533.

Sankararaman, S., S. Mallick, M. Dannemann, K. Prüfer, J. Kelso et al., 2014 The genomic landscape of Neanderthal ancestry in present-day humans. Nature 507: 354–357.

Schiffels, S., and R. Durbin, 2014 Inferring human population size and separation history from multiple genome sequences. Nat. Genet. 46: 919–925.

Schlötterer, C., R. Tobler, R. Kofler and V. Nolte, 2014 Sequencing pools of individuals—mining genome-wide polymorphism data without big funding. Nature Reviews Genetics 15: 749.

Schrider, D., J. Ayroles, D. R. Matute and A. D. Kern, 2018 Supervised machine learning reveals introgressed loci in the genomes of *Drosophila simulans* and *D. sechellia*. PLoS Genet. 14: e1007341.

Schrider, D. R., and A. D. Kern, 2015 Inferring selective constraint from population genomic data suggests recent regulatory turnover in the human brain. Genome Biol. Evol. 7: 3511–3528.

Schrider, D. R., and A. D. Kern, 2016 S/HIC: Robust Identification of Soft and Hard Sweeps Using Machine Learning. PLoS Genet. 12: e1005928.

Schrider, D. R., and A. D. Kern, 2017 Soft sweeps are the dominant mode of adaptation in the human genome. Mol. Biol. Evol. 34: 1863–1877.

Schrider, D. R., and A. D. Kern, 2018 Supervised Machine Learning for Population Genetics: A New Paradigm. Trends Genet. 34: 301–312.

Schrider, D. R., F. K. Mendes, M. W. Hahn and A. D. Kern, 2015 Soft shoulders ahead: spurious signatures of soft and partial selective sweeps result from linked hard sweeps. Genetics 200: 267–284.

Sheehan, S., and Y. S. Song, 2016 Deep learning for population genetic inference. PLoS Comput. Biol. 12: e1004845.

Simonsen, K. L., G. A. Churchill and C. F. Aquadro, 1995 Properties of statistical tests of neutrality for DNA polymorphism data. Genetics 141: 413–429.

Simonyan, K., and A. Zisserman, 2014 Very deep convolutional networks for large-scale image recognition. arXiv preprint arXiv:1409.1556.

Smith, J., G. Coop, M. Stephens and J. Novembre, 2018 Estimating time to the common ancestor for a beneficial allele. Mol. Biol. Evol.

Snoek, J., H. Larochelle and R. P. Adams, 2012 Practical bayesian optimization of machine learning algorithms, pp. 2951–2959 in Advances in neural information processing systems.

Sohn, K.-A., Z. Ghahramani and E. P. Xing, 2012 Robust estimation of local genetic ancestry in admixed populations using a nonparametric Bayesian approach. Genetics 191: 1295–1308.

Srivastava, N., G. Hinton, A. Krizhevsky, I. Sutskever and R. Salakhutdinov, 2014 Dropout: A simple way to prevent neural networks from overfitting. The Journal of Machine Learning Research 15: 1929–1958.

Sugden, L. A., E. G. Atkinson, A. P. Fischer, S. Rong, B. M. Henn et al., 2018 Localization of adaptive variants in human genomes using averaged one-dependence estimation. Nature Communications 9: 703.

Szegedy, C., W. Liu, Y. Jia, P. Sermanet, S. Reed et al., 2015 Going deeper with convolutions, pp. in CVPR.

Tajima, F., 1989 Statistical method for testing the neutral mutation hypothesis by DNA polymorphism. Genetics 123: 585–595.

Tennessen, J. A., A. W. Bigham, T. D. O’Connor, W. Fu, E. E. Kenny et al., 2012 Evolution and functional impact of rare coding variation from deep sequencing of human exomes. Science 337: 64–69.

Teshima, K. M., and H. Innan, 2009 mbs: modifying Hudson’s ms software to generate samples of DNA sequences with a biallelic site under selection. BMC Bioinformatics 10: 166.

Thornton, K. R., 2014 A C++ template library for efficient forward-time population genetic simulation of large populations. Genetics 198: 157–166.

Turner, T. L., M. W. Hahn and S. V. Nuzhdin, 2005 Genomic islands of speciation in Anopheles gambiae. PLoS Biol. 3: e285.

Voight, B. F., S. Kudaravalli, X. Wen and J. K. Pritchard, 2006 A map of recent positive selection in the human genome. PLoS Biol. 4: e72.

Vy, H. M. T., and Y. Kim, 2015 A Composite-Likelihood Method for Detecting Incomplete Selective Sweep from Population Genomic Data. Genetics 200: 633–649.

Washburn, J. D., M. K. M. Guerra, G. Ramstein, K. A. Kremling, R. Valluru et al., 2018 Evolutionarily informed deep learning methods: Predicting transcript abundance from DNA sequence. bioRxiv: 372367.

Yu, F., and V. Koltun, 2015 Multi-scale context aggregation by dilated convolutions. arXiv preprint arXiv:1511.07122.

Zaheer, M., S. Kottur, S. Ravanbakhsh, B. Poczos, R. R. Salakhutdinov et al., 2017 Deep sets, pp. 3394–3404 in Advances in Neural Information Processing Systems.

